# From protection to amplification: Imperfect chytridiomycosis prophylaxis increases infections in wild amphibians

**DOI:** 10.64898/2026.05.15.725113

**Authors:** K.M. Barnett, T.A. McMahon, A.D. Shepack, H.N. Buelow, Z. Barkley, A.V. Belsare, M. Risin, O. Milloway, J. Carozza, J. Beasley, B.K. Hobart, W.E. Moss, T. McDevitt-Galles, S. Detmering, B.A. Hilgendorff, C.L. Nordheim-Maestas, D.M. Calhoun, J.R. Rohr, P.T.J. Johnson, D.J. Civitello

**Affiliations:** Department of Biology, Emory University, 1510 Clifton Rd NE, Atlanta, Georgia 30322, USA; Department of Biology, Connecticut College, 270 Mohegan Ave Pkwy, New London, Connecticut 06320, USA; Department of Biology, University of Tampa, 401 W. Kennedy Blvd., Tampa, FL 33606, USA; Department of Biology, University of Notre Dame, Holy Cross Dr, Notre Dame, IN 46556, USA; Department of Ecology and Evolutionary Biology, University of Colorado, Boulder, 1900 Pleasant St, Boulder, CO 80309, USA; College of Forestry, Wildlife and Environment, Auburn University, Auburn, Alabama 36849, USA; Department of Forest and Wildlife Ecology, University of Wisconsin, Madison, WI 53706, USA; Department of Ecology, Evolution, and Marine Biology, University of California, Santa Barbara, CA, USA

**Keywords:** amphibian conservation, chytridiomycosis, imperfect vaccination

## Abstract

Wildlife vaccination could become a powerful strategy to mitigate disease-induced biodiversity losses, yet many vaccines for wildlife diseases provide only limited protection. Notably, tools to control the fungal pathogen *Batrachochytrium dendrobatidis* (Bd) are urgently needed for amphibian conservation. Laboratory experiments have demonstrated that prophylactic exposure to Bd metabolites increases host resistance, significantly reducing infection intensity in amphibians subsequently challenged with live Bd. Because Bd metabolites are non-infectious and applied topically, this treatment has potential to be administered to waterbodies to vaccinate and protect amphibians. We developed an agent-based model that indicated imperfect vaccination could reduce or amplify Bd infections at the population level, depending on degree of enhanced resistance or tolerance. Utilizing a Before-After-Control-Impact design with ten years of data, we conducted an ecosystem-level trial where we applied low levels of Bd metabolites or a sham control treatment to ponds in California and subsequently quantified Bd prevalence and infection intensity in metamorphosing Pacific chorus frogs (*Pseudacris regilla*). Unexpectedly, infection intensity was significantly greater in treated ponds relative to control ponds following metabolite addition. Additional model simulations indicated that this could occur via two mechanisms: (1) if treatment greatly increased tolerance alone or in combination with smaller increases in resistance, or (2) if a deleterious environmental interaction caused the treatment to increase susceptibility, rather than promote resistance. Future research is needed to determine whether tolerance or environmental factors drove heightened Bd infection intensities in this field trial to identify contexts in which this treatment can be used as a conservation tool.

**Significance statement:** Although wildlife vaccination is increasingly explored as a strategy to mitigate disease-induced population declines, many available vaccines provide limited protection, requiring careful consideration to design successful conservation campaigns. Here, we use both an eco-epidemiological model and field manipulation experiment to assess the effectiveness of an imperfect prophylactic treatment (akin to a prototype vaccine) for chytridiomycosis, a disease implicated in the massive decline of amphibian biodiversity worldwide. We unexpectedly found that prophylaxis-treated ponds had higher pathogen loads relative to control populations and models suggest this could result from enhanced tolerance or an adverse environmental interaction.

## Introduction

Wildlife vaccination is a promising conservation tool to mitigate the risk of disease-induced biodiversity loss (1–3) and a powerful public health intervention for the prevention of disease spillover to humans and livestock (4, 5). Although historically difficult to implement, the feasibility of wildlife vaccination has increased in recent years due to the growing availability of environmentally distributed vaccines, such as oral vaccine baits (6). The potential strength of vaccination for wildlife conservation lies in its ability to disrupt transmission as protection generated for vaccinated individuals often indirectly protects unvaccinated individuals, a phenomenon termed “herd immunity” (7).

Ideally, vaccines provide “perfect” (or sterilizing) protection, wherein all vaccinated individuals have lifelong resistance against infection. However, in practice, most vaccines provide only partial protection which wanes in efficacy over time (8, 9) and thus success of vaccination campaigns is highly sensitive to the degree to which vaccination provides imperfect immunity that boosts resistance (i.e., reduces infection establishment, decreases within-host pathogen replication, increases pathogen clearance, or reduces pathogen shedding), tolerance (i.e., reduces infection-induced mortality, also known as anti-toxin or anti-disease immunity), or both. Specifically, imperfect immunity that boosts resistance generally reduces transmission and protects populations, whereas immunity that boosts tolerance may lead to greater pathogen transmission by extending the duration of infectiousness, and can favor the selection of hypervirulent strains (10–13). The magnitude of these protective or deleterious effects on populations is mediated by the proportion of the population that is immunized (i.e., vaccine coverage) and the degree to which vaccination alters individuals’ immunity phenotypes.

The severity of the threat chytridiomycosis poses to amphibian conservation has prompted research into several novel disease control methods, including those based on vaccination, microbiome manipulation, and antifungal treatment (14–22). Chytridiomycosis, caused by the globally-distributed aquatic fungal parasite *Batrachochytrium dendrobatidis* (Bd), has contributed to at least 90 extinctions and can cause rapid die-offs via intensity-dependent mortality (23). The discovery that tadpoles (larvae), metamorphic frogs, and adults exhibit lower intensity Bd infections following topical exposure to a low concentration of Bd metabolites (non-infectious chemicals released by Bd in liquid culture) suggests the possibility of a vaccine for chytridiomycosis (15, 20, 21, 24). We currently refer to this Bd metabolite treatment as a prophylaxis rather than a vaccine because it is unknown whether the acquired resistance response is antibody-mediated in the species we studied, and a study by Siomoko et al. found that treatment with Bd metabolites is associated with increased presence of Bd-inhibitory bacteria (22). According to previous work in other disease systems, partial increases in host resistance can be sufficient to substantially reduce population-level impacts of disease (25). Additionally, this prophylaxis has important advantages: it is effective through indirect application and does not contain any infectious agents, therefore having strong potential for environmental distribution. However, this Bd metabolite intervention has never been tested for conservation-relevant impacts on the population scale or in a natural setting.

Here, we tested this novel intervention in waterbodies. First, we built an agent-based eco-epidemiological model to generate hypotheses for how limited protective efficacy and population coverage would affect key epidemiological and conservation endpoints, such as infection intensity, infection prevalence, host population size and spillover capacity. To do so, we considered four mechanistic representations of imperfect immunity: anti-infection resistance, anti-growth resistance, anti-transmission resistance, or anti-disease immunity (Table 1). We generated predictions for the field trial from simulations implementing one of the four immune mechanisms. Given the functional equivalency of a vaccine and prophylaxis as preventative treatments, we use the term vaccination in relation to our model to indicate the generalizability of its applications, while referring to the Bd metabolite treatment itself as a prophylaxis. We then tested these model-driven predictions using a Before–After–Control–Impact (BACI) experiment. In this waterbody-level experiment, we administered the prophylactic treatment for two years at the whole waterbody scale in replicated ponds in northern California during the breeding season and measured infection prevalence and intensity in post-metamorphic frogs 1–2 months later and compared these with eight years of baseline field data. Following the experiment, we conducted additional model simulations implementing the tolerance mechanism as well as simulated scenarios in which vaccination had an adverse, rather than protective effect, on infection-, growth-, or transmission phenotypes.

**Table 1.** Vaccines can provide four mechanisms of imperfect immunity. wherein Bd metabolite treatment 1) decreases probability of infection establishment upon pathogen exposure (anti-infection immunity), 2) increases pathogen clearance (anti-growth immunity), 3) decreases rate of pathogen shedding (anti-transmission immunity), or 4) increases the infection intensity threshold above which disease-induced mortality occurs (anti-disease immunity; i.e., tolerance).

| Type of immunity | Trait definition | Mechanism of resistance or tolerance? |
| --- | --- | --- |
| Anti-infection | Decreases probability of infection establishment upon pathogen exposure | Resistance |
| Anti-growth | Increases pathogen clearance | Resistance |
| Anti-transmission | Decreases rate of pathogen shedding | Resistance |
| Anti-disease | Increases the infection intensity threshold above which disease-induced mortality occurs | Tolerance |

## Results

### Bd–amphibian–vaccine model

#### Modes of protection

Infection intensities decreased in model scenarios where vaccination boosted resistance, whether it was by reducing infection establishment (Fig. 1a), reducing pathogen shedding (Fig. S1a), or increasing infection clearance (Fig. S1b). However, infection intensities increased when vaccination boosted tolerance (Fig. 1c). Prevalence only decreased at very high levels of coverage with high levels of boosted anti-infection (Fig. S2a), anti-transmission (Fig. S2b), and anti-growth resistance (Fig. S2c), and this reduction was weakest under scenarios of anti-growth resistance. Median infection prevalence in unvaccinated scenarios was already 100% under our parameterization, so model scenarios showed that enhanced tolerance did not change infection prevalence (Fig. S2d). Frog population size increased with increasing levels of coverage and resistance (Figs. 1b and S3), but the effect was less strong for tolerance as high levels of coverage and efficacy were needed to increase the population size by 20% as compared to untreated (Fig. 1d). Zoospore density, as a measure of spillover risk (i.e., high zoospore densities indicate greater risk of pathogen spillover), decreased with boosts to all forms of resistance (Fig. S4a-c). However, zoospore densities minimally increased when vaccination boosted tolerance (Fig. S4d). With increasing vaccine-induced anti-infection resistance and coverage, infection intensities decrease in unvaccinated frogs within the population (Fig. 2a) and a greater proportion of unvaccinated frogs survive (Fig. 2c), evidence of herd immunity. In contrast, with increasing vaccine-induced tolerance and coverage, infection intensities did not change for unvaccinated frogs within the population (Fig. 2b) and the proportion of the surviving unvaccinated population reduces (Fig. 2c).

**Figure 1.**
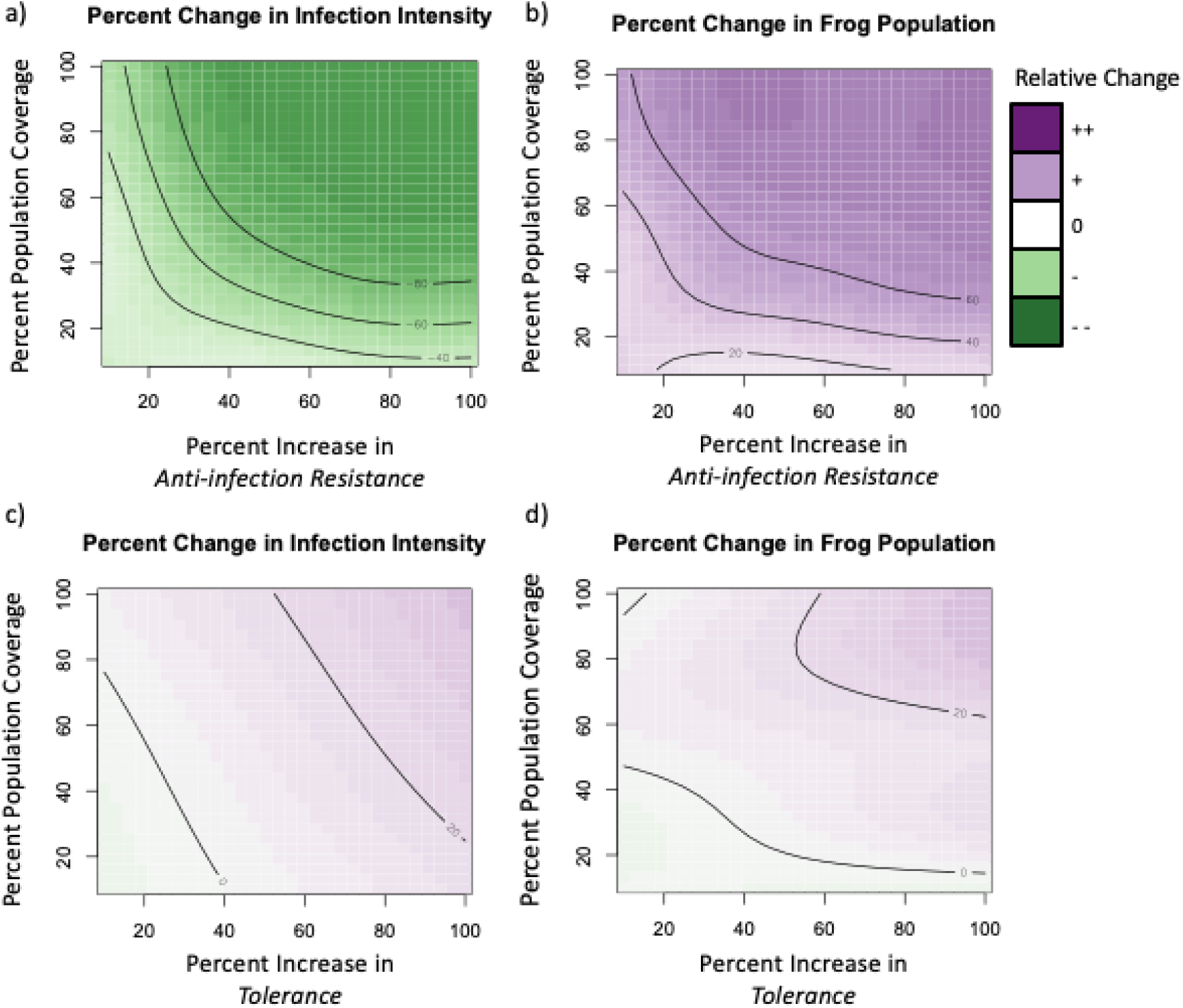
Percent change in infection intensity and frog population size when vaccination increases resistance or tolerance. Generalized Additive Model (GAM) summary of modeled changes in (a) infection intensity and (b) final frog population (green–purple color scale) as a function of increasing vaccination-induced anti-infection resistance (decrease in infection establishment; x-axis) and population coverage (y-axis), relative to simulations of an untreated control population. GAM summary of modeled changes in c) infection intensity and d) final frog population as vaccination increases host tolerance (increase in infection induced mortality threshold; x-axis) and population coverage on the y-axis relative to a modeled untreated control population. Deeper green shades represent reductions and deeper purples represent increases compared to unvaccinated populations. Contour lines define increments of 20% change relative to vaccine-free simulations. (a) Infection intensities decrease and, correspondingly, (b) frog population sizes increase as population coverage and efficacy of vaccine-induced resistance increases. Alternatively, (c) infection intensities increase as population coverage and efficacy of vaccine-induced tolerance increase and (d) population size only substantially increases with high levels of population coverage and boosted tolerance.

**Figure 2.**
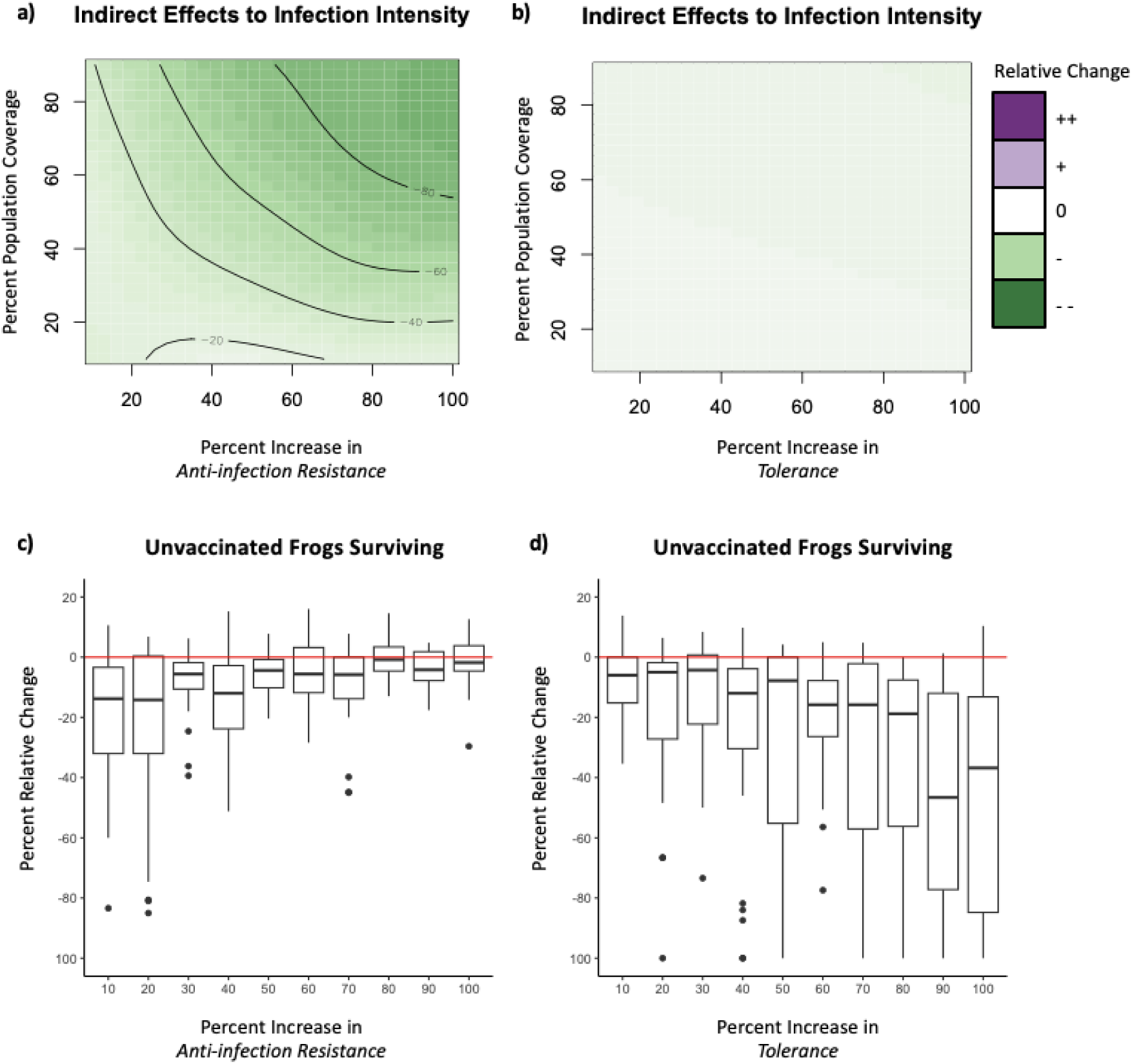
Indirect effects of vaccination on unvaccinated frog population. Generalized Additive Model (GAM) summary of modeled changes in infection intensity of unvaccinated surviving frogs (green–purple color scale) as a function of a) increasing vaccine-induced anti-infection resistance (decrease in infection establishment; x-axis) or b) vaccine-induced tolerance and population coverage (y-axis), relative to simulations of an untreated control population. Deeper green shades represent reductions and deeper purples represent increases in infection intensity as compared to unvaccinated populations. Contour lines define increments of 20% change relative to vaccine-free simulations. (a) Infection intensities decrease as population coverage and efficacy of vaccine-induced anti-infection resistance increases. In contrast, (b) infection intensities of surviving unvaccinated frogs do not change regardless of population coverage and efficacy of vaccine-induced tolerance. Percent relative change in the proportion of surviving frogs that are unvaccinated (non-negative values represent a protective effect i.e. herd immunity, wherein negative values represent harm to unvaccinated individuals) under different levels of vaccine-induced c) anti-infection resistance (x-axis) or d) tolerance (x-axis) in model scenarios with 50% population coverage. The red line at 0% indicates no change from the initial 50% unvaccinated composition. (c) As efficacy of vaccine-induced anti-infection resistance increases, more unvaccinated frogs survive to the end of the model run, indicating herd immunity. (d) As efficacy of vaccine-induced tolerance increases, fewer unvaccinated frogs survive to the end of the model run. Each boxplot represents the distribution of percent relative change (y-axis), which is the difference in the proportion of unvaccinated frogs surviving at the end of each model run compared to the initial 50% unvaccinated population, for a given increase in vaccine-induced a) anti-infection resistance (x-axis) or b) tolerance (x-axis).

### Field trial and challenge experiment

Bd infection intensity (i.e., mean Bd load of infected metamorphic frogs) and prevalence (i.e., proportion of population infected with Bd) were much lower in post-intervention years than in pre-intervention years for both control and treatment ponds. Control ponds showed an 88% decrease in estimated marginal mean infection intensity from pre- to post-intervention years (1,625, 95% CI 329-8,034 before intervention vs. 194, 95% CI 10-3,666 after intervention). In contrast, frogs from ponds that received Bd metabolite treatment had only a 23% decrease in estimated marginal mean infection intensity from pre- to post-intervention years (866, 95% CI 167-4,480 before intervention vs. 667, 95% CI 36-12,233 after intervention). If Bd metabolites had no effect on infection intensity, we would have expected intensities to decrease by a similar proportion from pre-intervention to post-intervention years. Instead, we found that infection intensities in treatment ponds decreased significantly less than those of frogs in control ponds. This time-by-treatment interaction (i.e., before vs. after intervention) was significant (P = 0.001; Fig. 3), and Bd infection intensities in treated ponds were 6.42 times greater than the expected value had infection intensities in treated ponds decreased similarly to those of control ponds.

**Figure 3.**
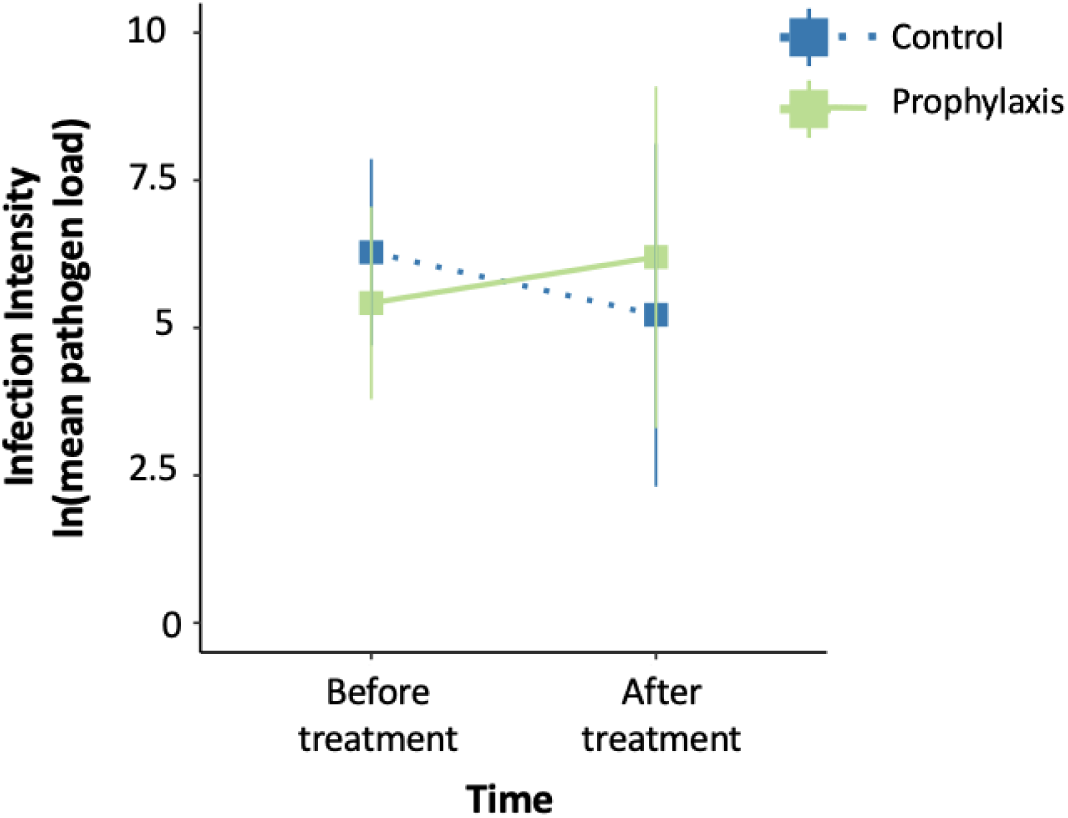
Estimated mean infection intensity (i.e., Bd load of infected individuals transformed to natural log scale) before and after Bd metabolite addition in *Pseudacris regilla* metamorphic frogs. There was a significant time by treatment interaction (p = 0.001) wherein frogs from ponds treated with Bd metabolites had significantly higher Bd loads after treatment than frogs in ponds treated with the sham treatment.

Control ponds had a 70% decrease in Bd infection prevalence from pre-intervention to post-intervention years (before intervention: 36%, 95% CI 0-83% vs. after intervention: 11%, 95% CI 6-82%). Treatment ponds had a similar 51% decrease in prevalence from pre- to post-intervention years (before intervention: 43%, 95% CI 8-87% vs. after intervention: 21%, 95% CI 1-91%). We found no significant time-by-treatment interaction in Bd infection prevalence for field-swabbed frogs (Fig. S5).

Among frogs challenged with live Bd in the laboratory, we found no significant time-by-treatment interaction in Bd infection intensity (Fig. S6b) or Bd infection prevalence (Fig. S6a). Infection intensities increased from pre- to post-intervention years in frogs from both control and treatment ponds. Lab-challenged frogs from control ponds had a final infection intensity of 8,333 (95% CI 3,150-22,042) Bd GE in the pre-intervention year and 20,674 (95% CI 8,066-52,989) Bd GE in the post-intervention year, amounting to a 148% increase from pre-intervention to post-intervention. Lab-challenged frogs from treatment ponds had a final infection intensity of 18,333 (95% CI 6,718-50,028) Bd GE in the pre-intervention year and 37,841 (95% CI 14,932-95,896) Bd GE in the post-intervention year, increasing by 106% between pre-intervention and post-intervention years. Infection prevalence in lab-challenged frogs increased similarly from pre-intervention to post-intervention years in control and treatment ponds.

### Additional Model Simulations

#### Two-way interactions with tolerance

When vaccination enhances both tolerance and resistance, the effect of tolerance on increasing Bd infection intensities was counteracted with increasing efficacy of the boosted resistance phenotype (Figs. 4a and S7). When combined with boosted tolerance, prevalence only decreased with a high degree of anti-infection resistance, otherwise, prevalence remained unchanged (Fig. S8). Host population sizes increased with increasing resistance (Figs. 4b and S9). While resistance phenotypes appeared to drive this boost in population size (Figs. 4b and S9), there appeared to be a slight observable interaction wherein boosts to tolerance increased population sizes when vaccination only provided weak anti-growth resistance (Fig. 4b). Lastly, zoospore densities decreased with increasing resistance efficacy and zoospore densities were slightly lower when tolerance was also low (Fig. S10).

**Figure 4.**
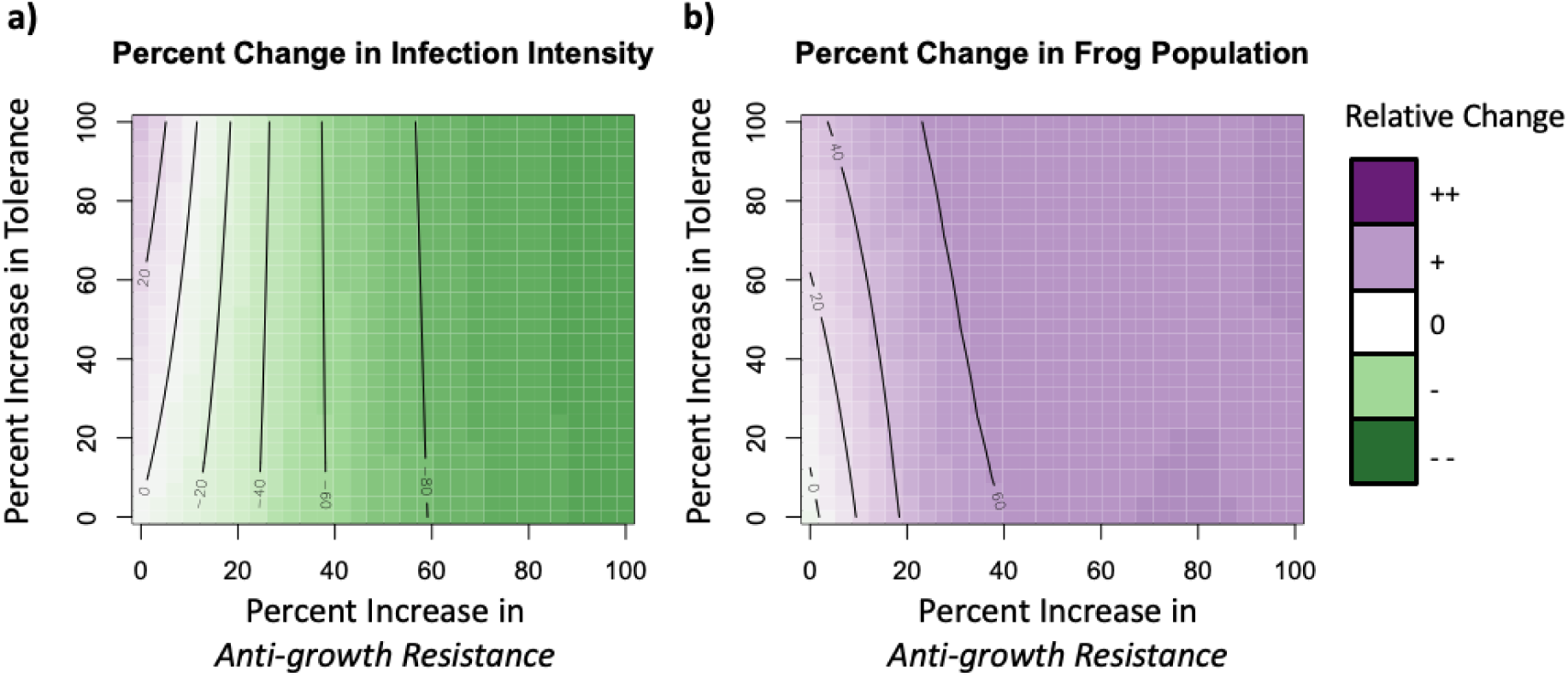
Percent changes to infection intensity and frog population size when vaccination provides both tolerance and resistance. Generalized Additive Model summary of modeled changes in a) infection intensity and b) final frog population (green–purple color scale) as vaccination boosts both anti-growth resistance (increase in pathogen clearance; x-axis) and tolerance (increase in infection induced mortality threshold; y-axis) in a population where 75% of hosts are treated, relative to simulations of an untreated control population. Deeper green shades represent reductions and deeper purples represent increases compared to unvaccinated populations. Contour lines define increments of 20% change relative to vaccine-free simulations. (a) When vaccination strongly boosts tolerance and only provides a minor increase to anti-growth resistance, infection intensities increase but when vaccination provides at least a small boost in anti-growth resistance, regardless of the degree of enhanced tolerance, infection intensities decrease. (b) Frog population sizes increase with increasing anti-growth resistance, with a minor boost from increasing tolerance at low levels of anti-growth resistance.

#### Hypothetical adverse reactions to vaccination

When we modeled scenarios where vaccination causes adverse reactions, infection intensities increased with greater reductions in resistance (i.e., increasing infection susceptibility, pathogen growth, or pathogen shedding) and greater population coverage (Figs. 5a and S11a,b). However, in model scenarios where vaccination reduced tolerance, infection intensities decreased with increasing coverage (Fig. S11c). Infection prevalence only decreased under a small range of scenarios where vaccination decreased anti-infection resistance, anti-transmission resistance, or tolerance at high levels of population coverage (Fig. S12a,b,d). Infection prevalence remained unchanged across all scenarios of decreased anti-growth resistance (Fig. S12c). As expected, lowered resistance and tolerance led to smaller resulting population sizes as compared to untreated populations (Figs. 5b and S13). Lastly, zoospore densities increased in scenarios where vaccination reduced resistance (Fig. S14a-c). Conversely, zoospore densities decreased in scenarios where vaccination weakened tolerance (Fig. S14d).

**Figure 5.**
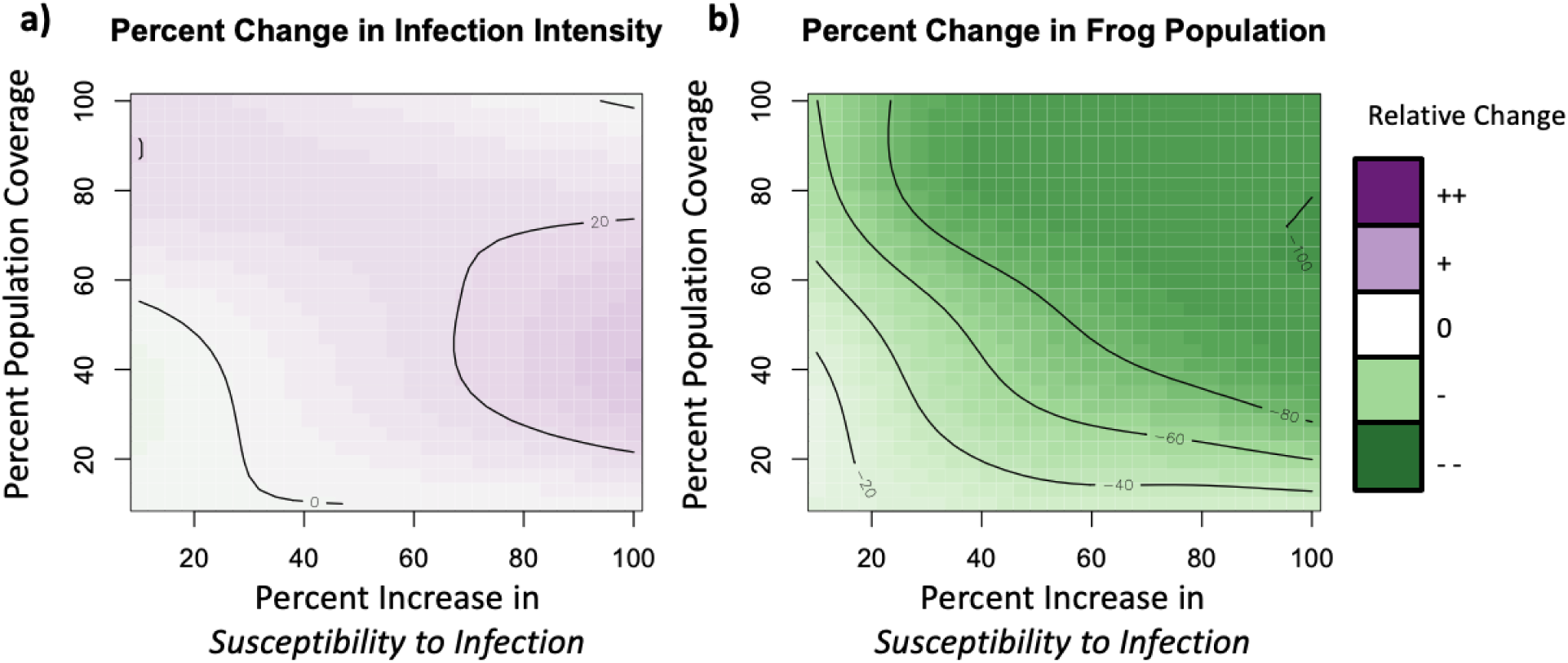
Changes to infection intensity and frog population size when vaccination is harmful. Generalized Additive Model summary of modeled changes in a) infection intensity and b) final frog population (green–purple color scale) as vaccination increases susceptibility to infection (increases infection establishment; x-axis) and population coverage (y-axis), relative to simulations of an untreated control population. Deeper green shades represent reductions and deeper purples represent increases compared to unvaccinated populations. Contour lines define increments of 20% change relative to vaccine-free simulations. (a) Infection intensities are higher in scenarios with moderate population coverage and high vaccine-induced susceptibility. b) Frog population sizes decrease as susceptibility and population coverage increase.

## Discussion

We conducted a field evaluation of a Bd prophylaxis that has previously been shown to induce resistance in amphibian hosts during laboratory trials, as indicated by a reduction in Bd infection intensities (15, 20, 21). General epidemiological models of vaccines as well as our amphibian-Bd specific agent-based model indicate that partial resistance via vaccination reduces infection intensity and prevalence in host populations (Figs. 1a, S1a-b, S2a-c). Counter to this hypothesis, we found that Bd infection intensity (i.e., Bd load of infected frogs) among prophylaxis-treated ponds was 6-fold higher than among sham-treated control sites (Fig. 3), even though all ponds exhibited an overall decrease in Bd infection over the study duration (likely due to drought or other climatic variables). This difference manifested only for infection intensity; we found no difference in infection prevalence between control and treatment ponds (Fig. S5). Among field-collected frogs challenged with Bd in the laboratory, there was no significant difference in infection intensity or probability of infection as a function of treatment (i.e., animals had comparable resistance; Fig. S6).

We used mechanistic modeling of amphibian–Bd–vaccine dynamics to complement our experiment and aid interpretation of these field results. All modeled scenarios where vaccination boosted a single resistance mechanism (e.g., reduced infection establishment, increased infection clearance, or reduced pathogen shedding) led to a decrease in infection loads (Figs. 1a and S1) and, at high levels of coverage and efficacy, a decrease in infection prevalence (Fig. S2). However, simulations in which vaccination increased only tolerance led to increases in infection intensity (Fig. 1c).

One potential explanation for our results is that prophylaxis greatly enhances tolerance, allowing treated frogs to sustain higher Bd infection loads prior to succumbing to disease-induced mortality. This modeling inference is based on simulating a tolerance effect in the absence of any resistance effects for vaccination. However, it is inconsistent with our findings from several laboratory trials that demonstrated Bd metabolites induce resistance, though those studies did not evaluate tolerance (15, 20, 21, 24). Therefore, we simulated additional scenarios in which vaccination simultaneously increased tolerance and one of the three resistance mechanisms. In scenarios where vaccination provided a weak boost to resistance but a stronger boost to tolerance, infection intensities were higher than unvaccinated scenarios (Figs. 4a and S7). Therefore, it is possible that Bd metabolites provide both resistance, as observed in previous lab trials, and tolerance, but that the boost to tolerance is stronger than the boost to resistance. A final, alternative explanation is that an environmental interaction caused field administration of Bd metabolites to be deleterious (i.e., adverse immune reaction), which could also result in higher infection intensities (Figs. 5a and S11a-b).

Both of these explanations pose risks for conservation-motivated disease control. Increasing infection susceptibility reduces population size, countering the intervention’s objective (Figs. 5b and S13). Meanwhile, although tolerance is beneficial at the individual level and can even increase population sizes with high efficacy and population coverage, it can be deleterious at the population-level when, rather than dying, highly infected individuals survive and are able to continue transmitting for prolonged durations (10, 26). Though minimal, increases in zoospore (i.e., the infectious stage of the pathogen) density as a consequence of enabling higher infection intensities may increase risk of spillover to susceptible sympatric hosts (Fig. S4d) and our model found that as efficacy of tolerance-enhancing vaccination increased, fewer unvaccinated individuals survived (Fig. 2d).

Future research is needed to uncover the underlying mechanism driving this conservation-relevant outcome of Bd metabolite addition increasing infection intensities in California ponds. It is critical to assess whether environmental factors such as temperature, sunlight, water chemistry, alternative circulating Bd strains, transmission seasonality, pond size, hydroperiod, sympatric non-amphibian Bd hosts and tadpole density impact treatment efficacy, as many of these factors have already been found to influence Bd infection prevalence and environmental persistence of zoospores (27–30). If an environmental interaction caused the prophylaxis treatment to have counterintuitive results within California ponds, it is possible Bd metabolite addition may be suitable in other amphibian communities given that Bd is found across diverse ecosystems. Moreover, vaccine-induced changes to host tolerance are often overlooked when assessing vaccine efficacy and our results underscore the importance of considering the degree to which immunization affects transmission via modes of resistance and tolerance. We suggest more focus be placed on directly quantifying differences in pathogen transmission by measuring duration of transmission period and productivity of pathogen shedding in immunized individuals compared to untreated individuals to better project the success of vaccination or prophylaxis in reducing infection prevalence and burdens at the population-level. Our model results suggest that the adverse impacts of enhanced tolerance on increasing infection loads can be counteracted with increased resistance (Figs. 4a and S7). Thus, if direct evidence is found that prophylaxis enhances tolerance (e.g., reduces Bd-associated immunopathology), it may be considered for use in combination with other interventions which strongly boost resistance.

The duration of protection is also a key factor in determining vaccination success. While our challenge experiment did not find evidence that metabolite exposure during the larval period induced changes in resistance that endured past metamorphosis, our laboratory challenge experiment with field-collected frogs was limited by sample size (number of frogs collected per pond and number of years replicated) and our inability to confirm that the frogs collected from Bd metabolite treated ponds were sufficiently exposed to those metabolites. Thus, future controlled laboratory studies should investigate if and to what degree protection provided by Bd metabolite exposure wanes through time and development. Even if induced resistance is not maintained through metamorphosis, infections accumulated as a tadpole persist through metamorphosis (31). Metamorphosis is thought to be the most vulnerable stage for Bd-induced, load-dependent mortality (32), so heightened resistance only as a tadpole may still benefit metamorphic frogs reaching metamorphosis with lower infection burdens. Additionally, this model can be adapted in the future to explore scenarios that vary the duration of acquired protection. Furthermore, studies should evaluate potential non-target impacts of Bd metabolite addition on other species in these aquatic communities, including invertebrates, some of which have exhibited pathology in response to extremely high doses of Bd metabolites (33, 34).

Overall, our findings emphasize that determining the effectiveness of a partially protective vaccine or prophylaxis requires a cross-scale (individual to population-level) approach, concern for environmental factors which may alter vaccine efficacy, and specific attention to the degree to which vaccination affects transmission. Importantly, while we were not able to confirm which, if any, of the frogs we sampled from treated ponds had been exposed to metabolites, our field trial and model results provide insight into how prophylaxis alters infection dynamics at the population level, including indirect effects on untreated hosts that are not captured in individual-level laboratory trials but are central to evaluating conservation outcomes. Tools that boost resistance and prevent Bd from causing large-scale die-offs are a priority for amphibian conservation. While research supporting the use of Bd metabolites to induce acquired resistance in laboratory settings is strong (15, 20, 21, 24), more work is needed to evaluate the environmental conditions and immune mechanisms underpinning the safety and efficacy of its use in field environments.

## Materials and Methods

### Bd–amphibian–vaccine model

We built a stochastic, stage-structured, and spatial agent-based model (ABM) of Bd-amphibian dynamics to assess pathogen and host population-level outcomes under different vaccination efficacies, coverage levels (i.e., proportion of the population immunized), and modes of imperfect immunity using NetLogo Version 6.3.0 (35). The model contained density-dependent transmission via a free-living Bd zoospore stage and infection-induced mortality elements like that of a prior non-spatial Bd-amphibian ABM (36), and added spatial structure, within-pond movement, host and pathogen development, and functional representations of imperfect vaccination (13). The model simulated within-season dynamics of a single species starting with tadpoles that, conditional on survival, transitioned into metamorphs by the end of the simulation. We used discrete daily time steps, spanning from 0–90 days to represent the aquatic Bd transmission season of ephemeral ponds in California. Each simulation began with 0.2% of tadpoles infected with 100 zoosporangia (the mature reproductive stage of Bd).

Spatially, the model contained three types of environmental patches: 1) perimeter pond patches, 2) deep pond patches, and 3) terrestrial patches (Fig. S15). In the ponds included in our study, high densities of tadpoles are observed along the shoreline of ponds where the water is shallow, so we assumed that perimeter pond patches are hotspots of contact with pathogens whereas, while tadpoles can shed zoospores in neighboring deep pond areas, those zoospores are unlikely to contact hosts given the lower density of tadpoles per volume of water. Thus, in the model, tadpoles could move between perimeter pond patches but deposited zoospores to both perimeter and neighboring deep pond patches, while metamorphs moved between land and perimeter pond patches. Given that tadpoles in the model did not move to deep pond patches and Bd is an aquatic fungal pathogen, zoospores deposited to deep pond or terrestrial patches did not contribute to onward infection.

The major processes of the model can be split into amphibian phenology and ecology, implementation of acquired immunity (vaccination), between-host transmission, and within-host infection processes (Table S1). The model tracked host survival, infection status, zoosporangium load, and the environmental zoospore density in the water body through time to obtain relevant population- and ecosystem-level outcomes such as final population size, infection prevalence, average infection intensity, and spillover risk. Parameters were selected based on values from the literature or were selected to match appropriate phenological and infection patterns (Table S1).

Tadpoles could die with a daily chance of 6% (‘tad-mort’; day^-1^) and 25% of tadpoles moved among perimeter pond patches each day. They developed into metamorphs starting on day 55 with a daily probability of 11% and all tadpoles remaining on day 74 transitioned to metamorphs. Metamorphs could die with a baseline probability of 2% (‘meta-mort’; day^-1^) at each time step and moved between patches daily with 10% of the population on a land patch and 90% on perimeter pond patches. Zoospores could be removed from the environment through contact with a host via an exposure parameter (i.e., amount of environmental units each host is exposed to) of 0.25 or by background death rate of 2 day^-1^ (28). Upon contact with a host, each zoospore infected the host with a baseline establishment probability of 0.25 (‘est’; unitless). Successful zoospores developed into reproductive zoosporangia over a fixed 4-day period (37). Depending on the life stage of the host (i.e., tadpole or metamorph), vaccination status, the mode of vaccine protection and efficacy, and history of exposure to zoospores, frogs could vary in zoosporangia load (‘Spn’). Infected amphibians, those with a zoosporangia load > 0, shed zoospores into the environment with a baseline rate of 17.8 zoospores per zoosporangium per day and can clear zoosporangia with a baseline probability of 0.2 per day (36). Metamorphs retained immune traits, infection status, and zoosporangia load from their tadpole state (31) and could die due to Bd infection if their zoosporangia load equals or surpasses the maximum threshold (‘smax’). Depending on the mode of vaccine protection, the zoospore shedding rate, zoospore establishment probability, zoosporangium clearance probability, or Bd-induced mortality threshold of a vaccinated frog may differ from baseline values proportional to the vaccine’s efficacy. Using this model, we ran several scenarios and display the model results as contour plots created in R statistical software, version 4.5.2 (38) using generalized additive models (GAM) with a gaussian distribution (package: mgcv, function: fvisgam).

#### Modes of protection

Vaccine-induced immunity can provide four types of functional protective phenotypes (i.e., “modes of protection”) by 1) reducing successful infection establishment (anti-infection resistance), 2) reducing pathogen shedding (anti-transmission resistance), 3) increasing infection clearance (anti-growth resistance), or 4) increasing a host’s ability to survive infection (tolerance). First, we ran our model where vaccination modulates one mode of protection over varying vaccine coverages (i.e., proportion of the population vaccinated). In these scenarios, vaccination reduces infection establishment, decreases pathogen shedding, increases infection clearance, or boosts the threshold for infection-induced mortality by a 10–100% change to the baseline immune parameter in increments of 10. The degree of change to the baseline parameter is defined as efficacy. Vaccine coverage also varied 10–100% in increments of 10 across these scenarios. Each combination of mode of immunity, level of efficacy, and vaccine coverage was replicated 25 times for a total of 10,000 runs across the experiment. Additionally, we ran a baseline control scenario without vaccination 250 times. We compared outcomes from the varying vaccination scenarios to outputs of this control scenario to calculate relative differences to evaluate hypothetical intervention success (defined as an increase to population size and reduction in infection intensity, infection prevalence, and zoospore density).

#### Two-way interactions with tolerance

To investigate the possibility that vaccination simultaneously affected tolerance and a mode of resistance, we ran two-way interaction scenarios where vaccination increases tolerance by 0–100% in increments of 10, in conjunction with also either decreasing infection establishment, reducing shedding, or increasing zoosporangia clearance from 0–100% in increments of 10. Vaccination coverage remains constant at 75% across these simulations. Each tolerance by mode of resistance combination was run 25 times for a total of 9,075 runs across the experiment. We then compared these scenario outputs to the control scenarios as described (see *Modes of protection*).

#### Hypothetical adverse reactions to vaccination

Given the unexpected outcome of the field experiment, we modeled the following scenarios: vaccination increases infection establishment, increases zoosporangia shedding, decreases tolerance or reduces infection clearance by 10–100% in increments of 10. We ran these scenarios across a gradient of coverages ranging from 10–100% in increments of 10. Again, each scenario was replicated 25 times for a total of 10,000 runs across the experiment and outputs were compared to results from an unvaccinated control scenario.

### Field trial

#### Experimental design

We used a replicated, whole-waterbody Before–After–Control–Impact (BACI) experimental design to test the efficacy of an environmentally administered Bd metabolite prophylaxis in *Pseudacris regilla* (Pacific chorus frog) populations. Experimental units were ponds in the Blue Oak Ranch UC Reserve and Joseph D. Grant County Park in Santa Clara County, California, USA (Permit #14025, 19-940383, 21-1194611, and 1389361). We chose *P. regilla* as the focal species for this study as they have been implicated as a reservoir species for *Batrachochytrium dendrobatidis* (26), have stable populations (39), and laboratory experiments found that exposure to Bd metabolites can induce resistance in *P. regilla* tadpoles and metamorphs during subsequent challenge with live Bd (21, 24). Moreover, *P. regilla* has been identified as the most important host for Bd persistence in the ponds we study given its ability to maintain Bd infections and wide dispersal (40). We collected pre-intervention baseline data on pond-level Bd prevalence and load between 2011 and 2019 (2020 data unavailable due to COVID-19 pandemic restrictions); ponds varied in the number of years with pre-intervention data available, but all had a minimum of two years of data included (mean duration 4.6 years, range = 2 – 8 years). In pre-intervention years 2011 to 2018, we swabbed an average of 9 (min: 1, max: 15) emerging metamorphs (Gosner Stage 44–46; hereon, referred to as both “metamorphs” or “frogs”) per year for each pond. We increased our pre-intervention swabbing effort in 2019, swabbing an average of 36 (min: 25, max: 48) metamorphs per pond. In the springs of 2021 and 2022, we dosed 6 ponds with Bd metabolites (3000 zoospores removed/L pondwater per dose) and 6 ponds with a sham control treatment (production described below) 4 times per year. Ponds were selected randomly across groups stratified by size, historical Bd prevalence and intensities, and amphibian community composition. Timing of treatment administration was chosen according to host phenology; we dosed ponds at peak tadpole density (mid to late April) which was approximated as the period after eggs hatched, before tadpoles metamorphosed and after breeding adults had retreated. In the summers of 2021 and 2022, we swabbed an average of 38 metamorphs (min: 7, max: 76) per pond per year. Nested within this study, we conducted a BACI-designed challenge experiment to quantify post-metamorphosis infection loads given exposure to a known live pathogen dose to evaluate if metamorphs from treated ponds had enhanced resistance. In 2019 (pre-intervention) and 2022 (post-intervention), we collected a subset (min: 10, max: 50, average: 33 per pond per year) of metamorphs from the field and dosed them with a known quantity of Bd to quantify pre- and post-intervention resistance. Additional methods for the live Bd challenge experiment can be found in the *Supporting Information* materials.

#### Preparing and administering Bd metabolites

We prepared the Bd metabolite stock following methods previously described in Nordheim et al. (15). To summarize, we flooded Bd+ agar plates with artificial spring water (ASW, 41) to obtain a solution containing live Bd and metabolites and then calculated the concentration of Bd zoospores in the solution using a 10 µL aliquot of the Bd+ solution on a hemocytometer to estimate the quantity of metabolites. Then, Bd zoospores and zoosporangia were removed from the solution by passing it through a 1.2 μm filter (GE Whatman Laboratory Products), thereby obtaining a filtrate containing only Bd metabolites suspended in ASW and no infectious material. To produce the sham control treatment, we replicated all steps for Bd metabolite stock preparation, with the exception of using Bd-, rather than Bd+, agar plates. The Bd metabolite stock and sham control were kept frozen until thawed prior to administration. Pond volume was determined to quantify the amount of Bd metabolite stock needed and was estimated using field measurements of perimeter, surface area, and depth at the pond center. We diluted the Bd metabolite stock into pond water accordingly to attain an overall pond-level concentration of approximately 3000 zoospores-removed per L for each dosing event. In keeping with past published work, we refer to the concentration of the treatment as zoospores removed per L in reference to the pre-filtration concentration of Bd zoospores (20–22). We used the average Bd metabolite stock concentration for the sham control dilution factor. Tadpoles were often observed congregating at the shoreline (personal observation), thus we used watering cans to distribute the diluted metabolite or sham treatment along the perimeter of each pond, administering metabolites from shoreline to approximately 1.5 m off the shore. We dosed each pond four times (2021 and 2022) over two weeks in April of both years.

#### Pond-level Bd infection prevalence and load

Due to the scale of the project, some factors such as swabbing technique, quantitative polymerase chain reaction (qPCR) protocol, and lab varied between years within this study, but all methods were kept standard within-year and uniform across treatment groups, thus being accounted for within the BACI design structure. For field swabs collected from 2011–2021, metamorphs were swabbed using MW113 swabs (Advantage Bundling, North Carolina, USA) on the underside of their head, ventral surface, vent, cloaca, legs, and arms 10× each per location (total of 70 swab strokes). Bd infection status and load on swabs were determined by qPCR (see 42) with plasmid standards designed to target Bd/Bsal (Pisces Molecular, Boulder, Colorado, USA). We screened for inhibition in every sample using TaqMan Exogenous Internal Control Reagents (Applied Biosystems, Foster City, California USA) and any sample with inhibition was rerun. In 2022, metamorphs were swabbed 10× on the ventral patch and 10× on each leg (a total of 30 strokes) with the same MW113 swabs and swabs were processed using the same methodology. In 2022, metamorphs were sent to the lab individually as part of the challenge experiment (see below) and the same swabbing and qPCR processing methods were used in the lab as in the field, and so the initial swabs taken upon arrival to the lab were used for the field swab dataset. Bd infection loads for swabs processed in 2011-2018 were reported in zoospore equivalents, whereas Bd infection loads were reported in genome equivalents for 2019, 2021, and 2022.

#### Data analysis on Bd field trial swabs

We conducted all statistical analyses in this study using R statistical software, version 4.5.2 (38). To assess if Bd metabolite addition altered infection prevalence in field-swabbed metamorphs, we used a binomial generalized linear model on binary (individual-level) infection status, with time (before or after intervention) crossed with treatment (sham or Bd metabolites) as predictors for Bd prevalence and year and pond as random effects (package: glmmTMB, function: glmmTMB). We conducted a likelihood ratio test (package: stats, function: anova) to evaluate significance against a null model. We calculated confidence intervals for Bd prevalence using the marginaleffects package (function: estimate_means). To test for a time × treatment interactive effect on infection intensity, we used a zero-inflated negative binomial generalized linear model (package: glmmTMB, function: glmmTMB) with raw Bd load per individual as the response, time × treatment as predictors, and pond and year as random effects and fitted zero-inflation with these covariates. The model is written bd_load ∼ treatment*before.after + (1 | pond) + (1 | year) wherein “before.after” refers timing of treatment relative to pre- or post-intervention, with the following predictors for both the conditional (count) component of the model and the zero-inflation component. We also used the marginaleffects package (function: estimate_means, estimate = “average”) to extract the marginal mean infection intensity estimates for each treatment.

### Laboratory Infection Challenge Experiment

As a subset of the Before–After–Control–Impact field trial, we conducted a live Bd challenge experiment to assess if there were differences in post-metamorphosis resistance between frogs from ponds treated with Bd metabolites or sham.

#### Pre-intervention

In 2019, *P. regilla* metamorphs (defined as Gosner Stage 44–46; n= 30–50 per pond, average = 40) from each pond were sent overnight on ice in groups of 15–25 frogs per 1000mL Tupperware container to the McMahon Lab at University of Tampa, Tampa, Florida, USA (IACUC #2018-2). Each container contained a moistened paper towel and air holes in the lid. On day of arrival, metamorphs were swabbed 10× on the left leg, weighed, and placed in 12 oz clear deli cups with air holes and paper towel dampened with artificial spring water (ASW, 41) on the bottom. They were kept on a 12 hr light/12 hr dark cycle at 21 degrees Celsius. Metamorphs were fed live calcium-dusted crickets 3× per week and container changes were done weekly. All metamorphs were dosed with 6 × 10^4^ zoospores of live Bd JEL-270 (isolated from California) and were swabbed, weighed and euthanized on the 10th day after Bd exposure using Orajel (20% benzocaine gel was placed on the head and dorsal side of the frog (43). Bd infection status and load were quantified using the same methods used for field swabs.

#### Post-intervention

In 2022, *Pseudacris regilla* metamorphs (n= 9–45 per pond, average = 28) were sent overnight on ice in individual falcon tubes, with a moistened cotton ball and air hole, to the Rohr Lab at University of Notre Dame, South Bend, Indiana, USA (IACUC #19-04-5328). Upon arrival, each metamorph was swabbed according to the protocol used for the 2022 field swabs and weighed. Metamorphs were kept in the same housing conditions as those in 2019. Metamorphs were split into two batches per arrival date – half of the frogs were challenged with 2.5 × 10^5^ zoospores of live JEL-270 Bd on day of arrival and the other half were challenged with the same dose of live Bd a week after to standardize effects of Bd batch. As in 2019, frogs were swabbed, weighed and euthanized on the 10th day after Bd exposure using Orajel (20% benzocaine gel was placed on the head and dorsal side of the frog;, 43). Bd infection status and load were diagnosed using the same methodology as that used for the pre-intervention challenge experiment swabs and field collected swabs.

#### Data analysis

To test if Bd metabolite addition altered infection outcomes in field-collected metamorphs challenged with a known dose of live Bd, we used a binomial generalized linear model on binary infection status (“bd_end”) of the post-challenge swab with time crossed with treatment as predictors for probability of Bd infection and pond as a random effect (package: glmmTMB, function: glmmTMB) in R statistical software, version 4.5.2 (38). The model is written as bd_end ∼ treatment*before.after + (1 | pond) wherein “before.after” refers timing of treatment relative to pre- or post-intervention with a binomial error distribution. We excluded all metamorphs with a positive initial swab (Bd GE > 0) from the data analysis, leaving an average of 23 metamorphs (min: 8, max: 39) per pond per year in the analysis. We again used a likelihood ratio test (package: stats, function: anova) to evaluate significance against a null model and calculated confidence intervals using the marginaleffects package. Then, we used a zero-inflated negative binomial generalized linear model (package: glmmTMB, function: glmmTMB) with time × treatment as predictors for infection intensity with pond as a random effect and fitted zero-inflation with these covariates. This model is written bd_load ∼ treatment*before.after + (1 | pond) with a negative binomial error distribution.

## Acknowledgements

We thank Drs. Katia Koelle, Jaap de Roode, and Sarah Bowden for providing feedback on this manuscript and Dr. Joyce Longcore for providing the Bd strains. We thank land managers Dr. Zac Harlow of Blue Oak Ranch Reserve and Jared Bond and Don Rocha of Joseph D. Grant County Park for access to field sites and Johnson Lab members for collecting swabs. We greatly thank Dr. Cherie Briggs for sharing historical Bd data. We also acknowledge the lives of the frogs used in this study, without which this study could not have been conducted. Funding that helped support the experiments or collection of historical data were provided by grants from the National Science Foundation (IOS-1755002, IOS-1754862, IOS-2113544, DEB 1149308, DEB 1754171, GRFP DGE-193797), National Science Foundation/National Institutes of Health Ecology and Evolution of Infectious Diseases program grants (R01GM1094995, R01GM135935), Section 6 Grant from California Department of Fish and Wildlife and United States Fish and Wildlife Service (P188010), the Infectious Disease Across Scales Training Program, Laney Graduate School, Connecticut College Department of Biology and Emory University Department of Biology.

## Author contributions

K.M.B., D.J.C., T.A.M., P.T.J.J., J.R.R., and A.D.S. conceived the experiments and K.M.B., D.J.C., and A.B. conceived of the model design. K.M.B., Z.B., M.R., O.M., J.C., J.B., B.K.H., W.M., T.M., S.E.D., D.M.C. and C.L.N-M. conducted the fieldwork. T.A.M., A.D.S., H.N.B., K.M.B., S.E.D and B.A.H. conducted the lab experiments, processed samples, and prepared the treatments. K.M.B. and D.J.C. conducted the analyses and wrote the manuscript. D.J.C., T.A.M., P.T.J.J., J.R.R., and K.M.B. provided funding.

## Competing interest statement

We declare no competing interests.

## Notes

### Competing Interest Statement

The authors have declared no competing interest.

### Summary of Updates

Small revisions have been made to include additional citations, update author affiliations, and improve certain phrasings.

