## Supplemental Data 1 for "From protection to amplification: Imperfect chytridiomycosis prophylaxis increases infections in wild amphibians"

### Figures

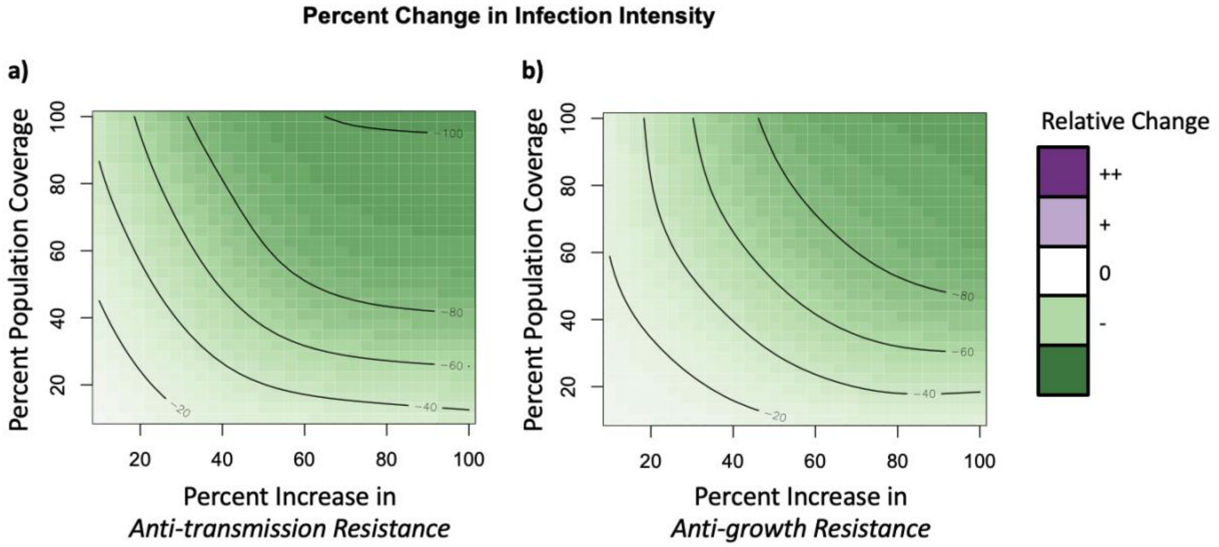


**Figure S1. Changes in infection intensity when vaccination provides either anti-transmission or anti-growth resistance across increasing population coverage.** Generalized Additive Model summary of modeled changes in infection intensity (green-purple color scale) as a function of vaccination-induced increase in (a) anti-transmission resistance (i.e., decrease in pathogen shedding) or (b) anti-growth resistance (increase in pathogen clearance; x-axis) and population coverage (y-axis), relative to simulations of an untreated control population. Deeper green shades represent reductions and deeper purples represent increases in infection intensity compared to unvaccinated populations. Contour lines define increments of 20% change relative to vaccine-free simulations. Infection intensities decrease as (a) anti-transmission resistance or (b) anti-growth resistance, as well as, population coverage increase.

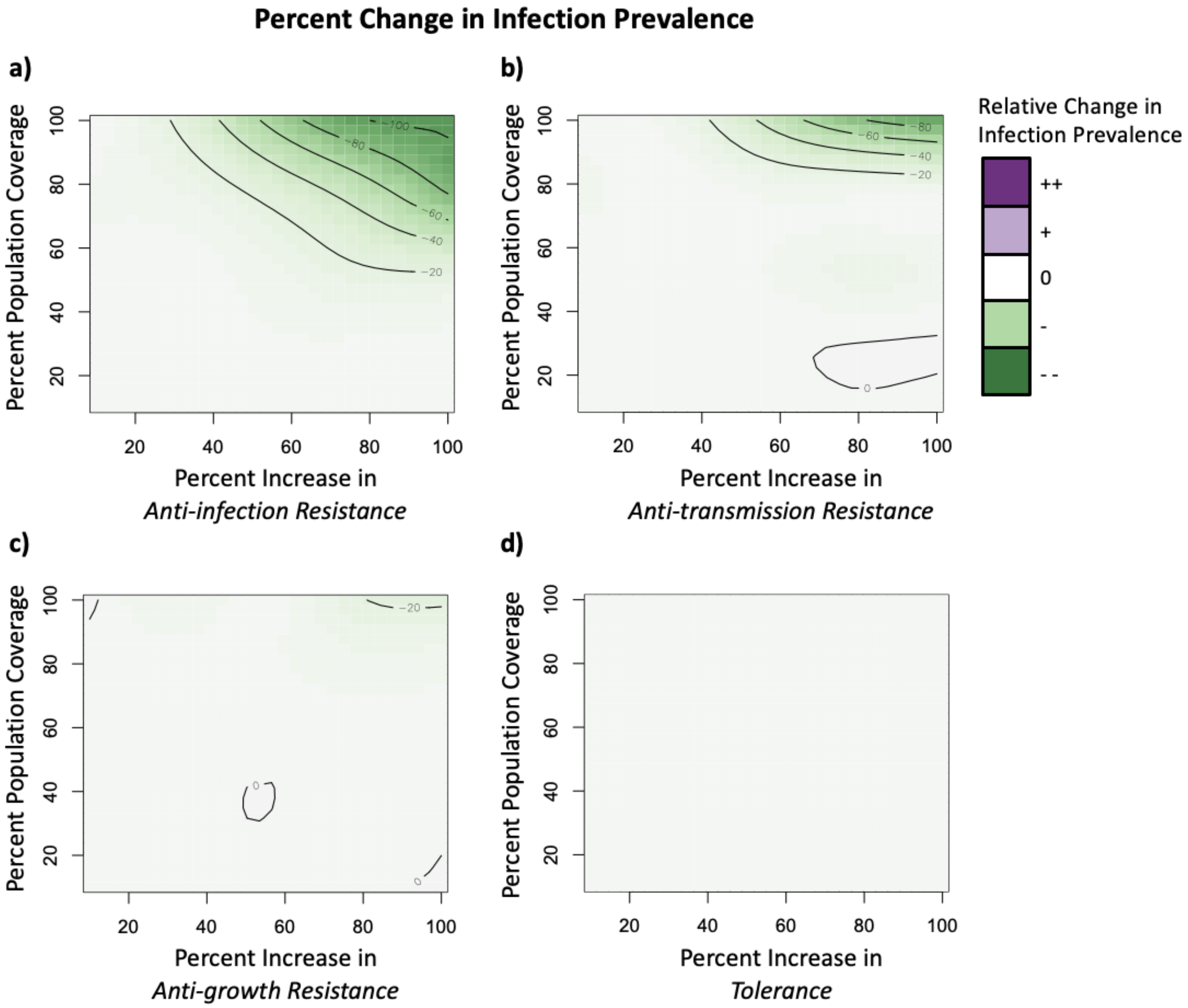


**Figure S2. Changes in infection prevalence when vaccination provides anti-infection resistance, anti-transmission resistance, anti-growth resistance or tolerance across increasing levels of population coverage.** Generalized Additive Model summary of modeled changes in infection prevalence (green-purple color scale) as a function of vaccination-induced increase in (a) anti-infection resistance (decrease in infection establishment), (b) anti-transmission resistance (decrease in pathogen shedding), (c) anti-growth resistance (increase in pathogen clearance), or (d) tolerance (increase in infection induced mortality threshold; x-axis) and population coverage (y-axis), relative to simulations of an untreated control population. Deeper green shades represent reductions and deeper purples represent increases in infection prevalence compared to unvaccinated populations. Contour lines define increments of 20% change relative to vaccine-free simulations. Infection prevalence decreases at high levels of (a) anti-infection resistance, (b) anti-transmission resistance, or (c) anti-growth resistance and coverage but does not change under any scenarios of (d) tolerance boosting vaccines.

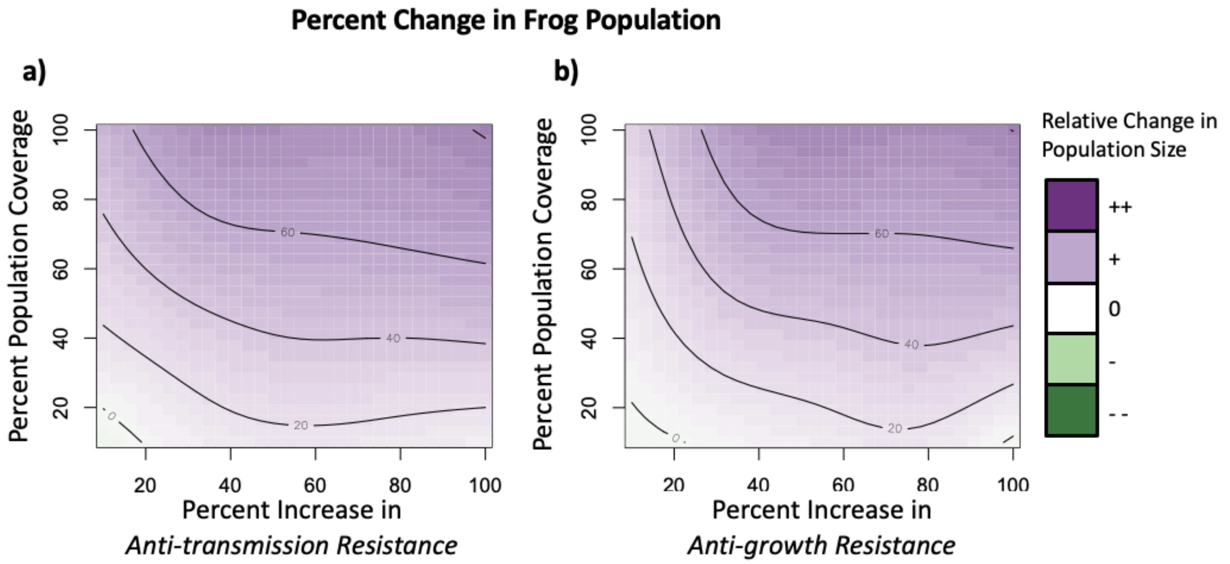


**Figure S3. Changes to frog population size when vaccination provides either anti-transmission or anti-growth resistance across increasing population coverage.** Generalized Additive Model summary of modeled changes in surviving population size (green-purple color scale) as a function of vaccination-induced increase in (a) anti-transmission resistance (i.e., decrease in pathogen shedding) or (b) anti-growth resistance (increase in pathogen clearance; x-axis) and population coverage (y-axis), relative to simulations of an untreated control population. Deeper green shades represent reductions and deeper purples represent increases in population size compared to unvaccinated populations. Contour lines define increments of 20% change relative to vaccine-free simulations. Frog population sizes increase as population coverage and (a) anti-transmission resistance or (b) anti-growth resistance increase.

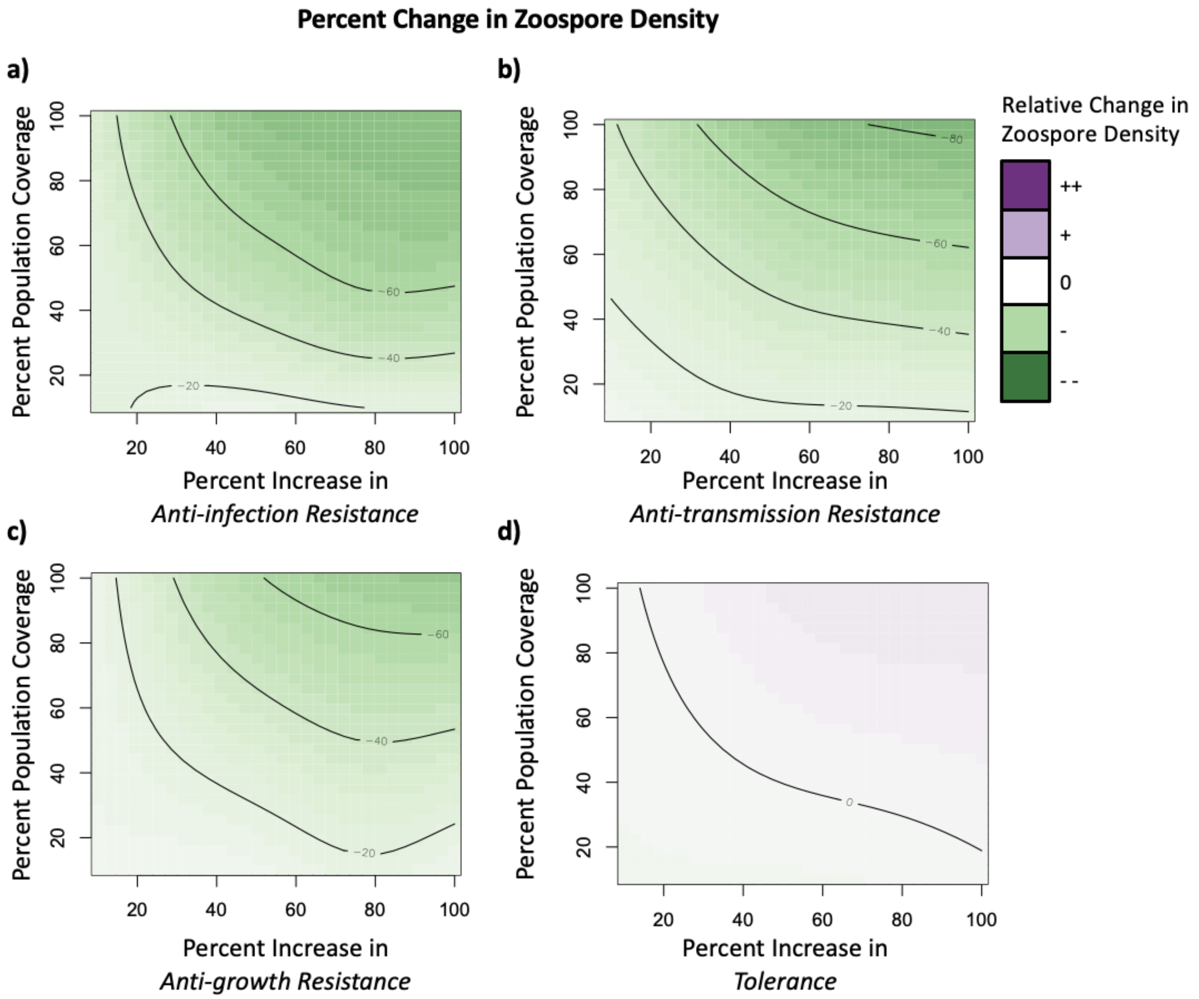


**Figure S4. Changes in zoospore density when vaccination provides anti-infection resistance, anti-transmission resistance, anti-growth resistance or tolerance across increasing levels of population coverage.** Generalized Additive Model summary of modeled changes in zoospore density (green-purple color scale) as a function of vaccination-induced increase in (a) anti-infection resistance (decrease in infection establishment), (b) anti-transmission resistance (decrease in pathogen shedding), (c) anti-growth resistance (increase in pathogen clearance), or (d) tolerance (increase in infection induced mortality threshold; x-axis) and population coverage (y-axis), relative to simulations of an untreated control population. Deeper green shades represent reductions and deeper purples represent increases in zoospore density compared to unvaccinated populations. Contour lines define increments of 20% change relative to vaccine-free simulations. Zoospore densities decrease as (a) anti-infection resistance, (b) anti-transmission resistance, or (c) anti-growth resistance and coverage increase but zoospore density remains unchanged with enhanced tolerance.

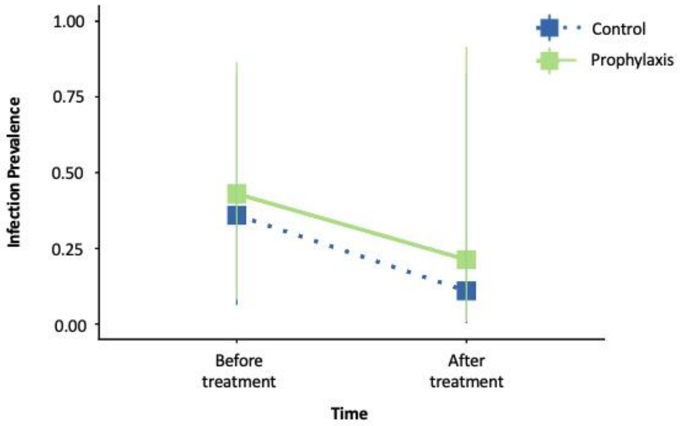


**Figure S5.** No significant interaction (P > 0.05) between time (before/after treatment addition) and treatment type for infection prevalence in field swabbed frogs.

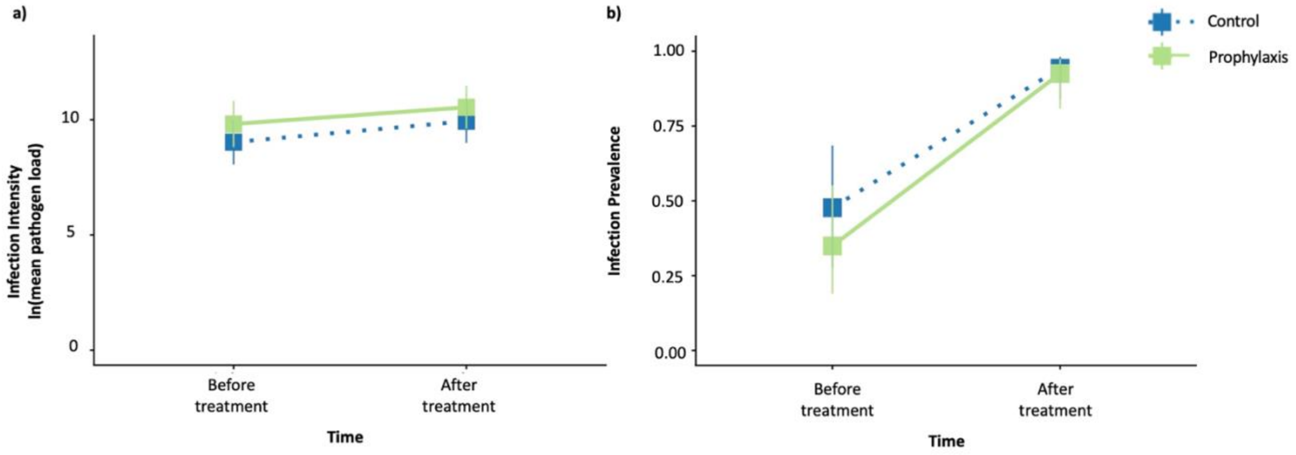


**Figure S6.** In the Bd challenge experiment, there was no significant interaction (P >0.05 for Fig S6 a and b) between time (before/after treatment addition) and treatment in a) estimated marginal mean infection intensity (Bd load of infected individuals transformed to natural log scale) or b) infection prevalence.

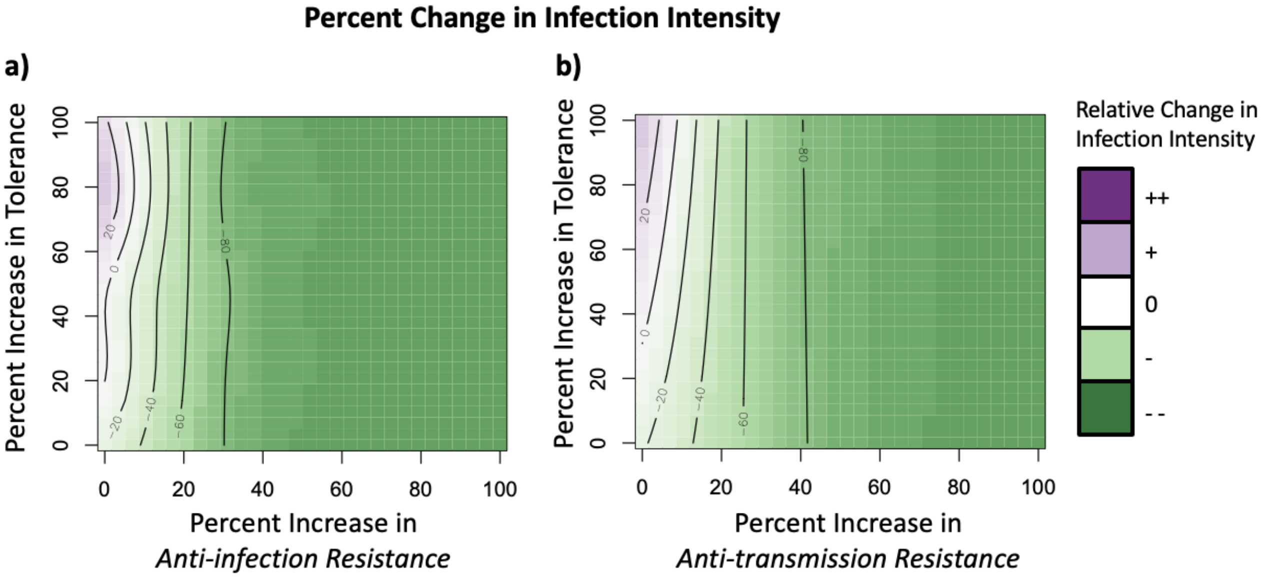


**Figure S7. Changes to infection intensity when vaccination provides both tolerance and either anti-infection or anti-transmission resistance.** Generalized Additive Model summary of modeled changes in infection intensity (green-purple color scale) as a function of vaccination-induced increases in (a) anti-infection resistance (decrease in infection establishment) or (b) anti-transmission resistance (decrease in pathogen shedding; x-axis) and enhanced tolerance (y-axis), relative to simulations of an untreated control population. Deeper green shades represent reductions and deeper purples represent increases in infection intensity as compared to unvaccinated populations. Contour lines define increments of 20% change relative to vaccine-free simulations. Infection intensities decrease as (a) anti-infection resistance or (b) anti-transmission resistance increase, but infection intensities increase at high levels of enhanced tolerance and low levels of resistance.

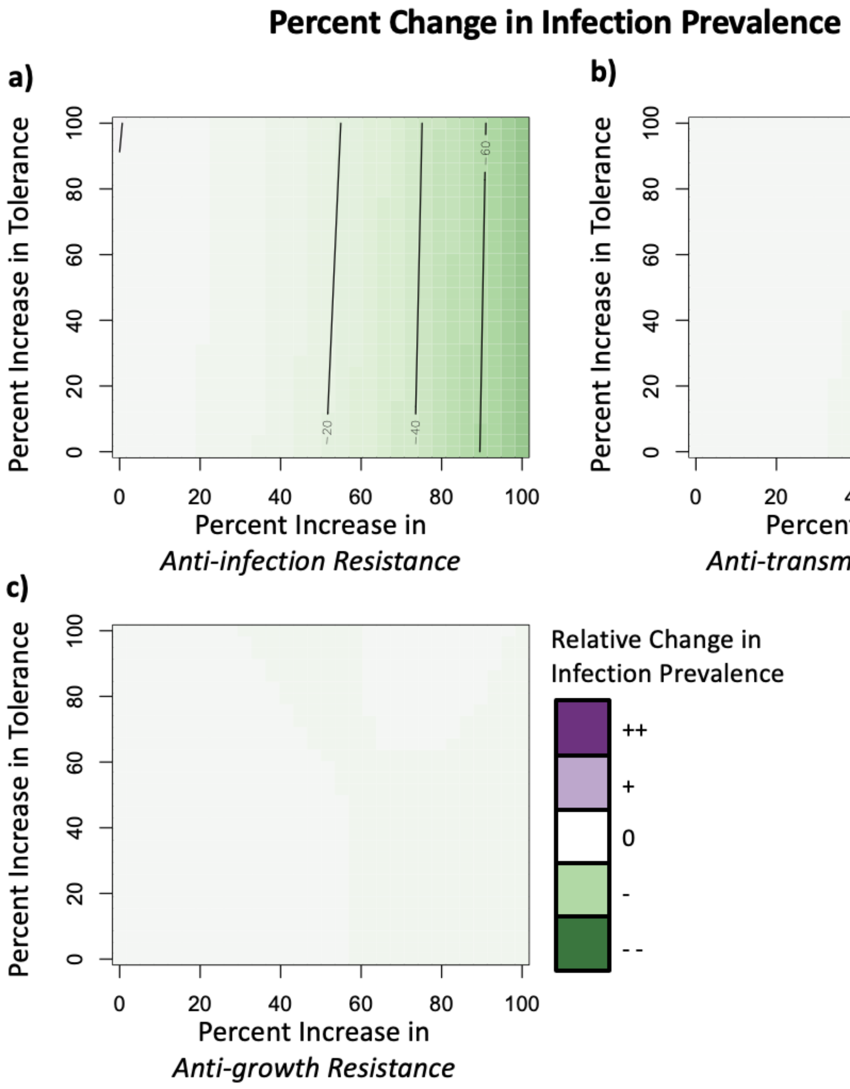


**Figure S8. Changes to infection prevalence when vaccination provides both tolerance and resistance.** Generalized Additive Model summary of modeled changes in infection prevalence (green-purple color scale) as a function of vaccination-induced increases in (a) anti-infection resistance (decrease in infection establishment), (b) anti-transmission resistance (decrease in pathogen shedding), (c) anti-growth resistance (increase in pathogen clearance; x-axis) and increase in tolerance (infection induced mortality threshold; y-axis) where 75% of hosts are vaccinated, relative to simulations of an untreated control population. Deeper green shades represent reductions and deeper purples represent increases in prevalence as compared to unvaccinated populations. Contour lines define increments of 20% change relative to vaccine-free simulations. Infection prevalence decreases at high levels of (a) anti-infection resistance, regardless of level of boosted tolerance, but does not change under any combinations of boosted (b) anti-transmission resistance or (c) anti-growth resistance and enhanced tolerance.

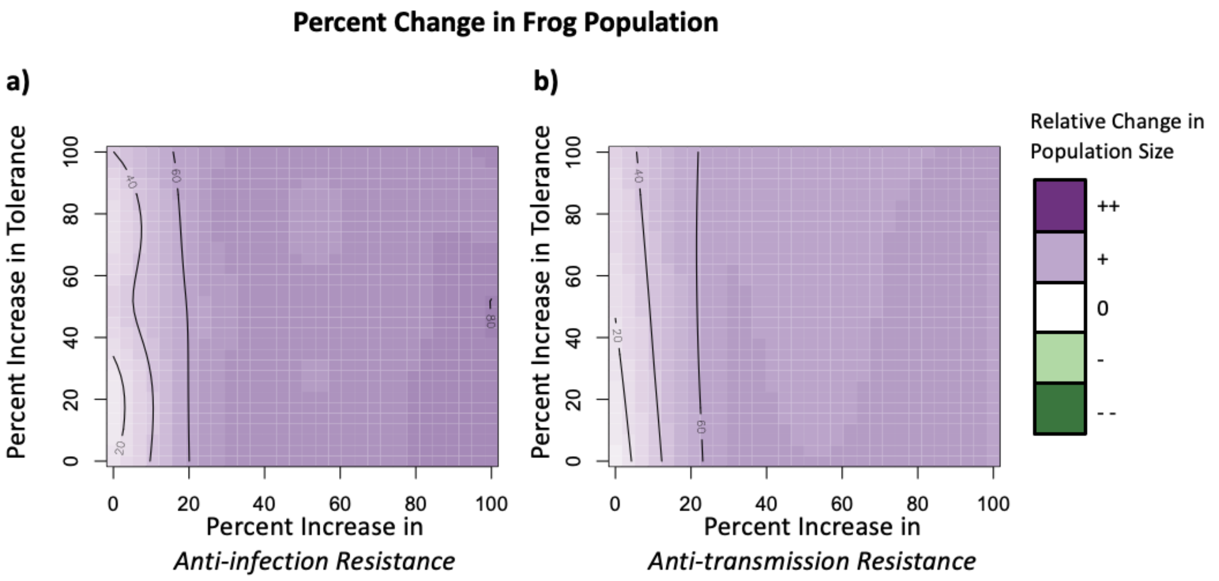


**Figure S9. Changes to frog population size when vaccination provides both tolerance and either anti-infection or anti-transmission resistance.** Generalized Additive Model summary of modeled changes in surviving frog population size (green-purple color scale) as a function of vaccination-induced increases in (a) anti-infection resistance (decrease in infection establishment) or (b) anti-transmission resistance; (decrease in pathogen shedding; x-axis) and enhanced tolerance (y-axis) in a population where 75% of hosts are treated, relative to simulations of an untreated control population. Deeper green shades represent reductions and deeper purples represent increases in population size as compared to unvaccinated populations. Contour lines define increments of 20% change relative to vaccine-free simulations. Population sizes increase as (a) anti-infection resistance or (b) anti-transmission resistance, with negligible effects of increasing tolerance.

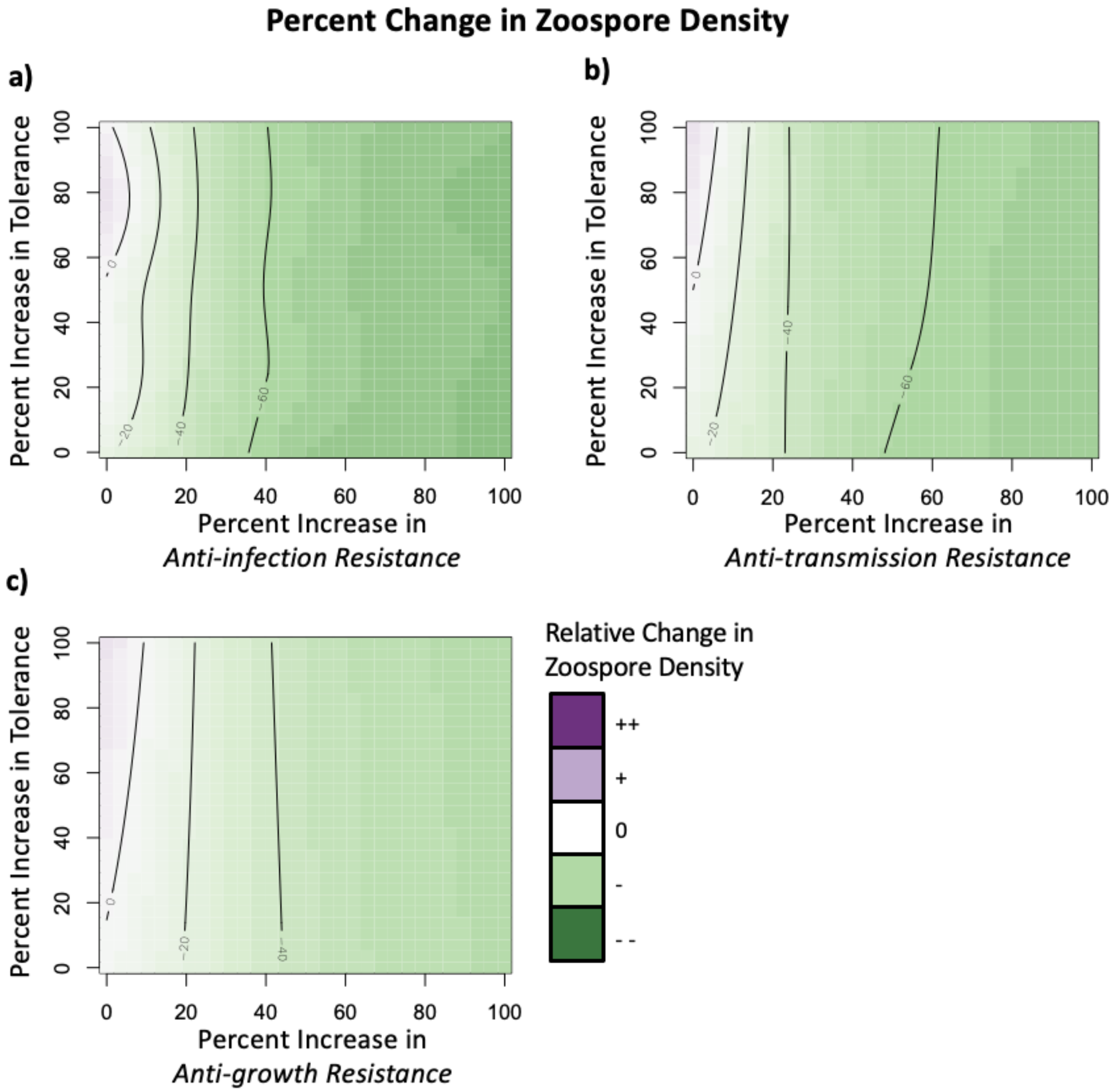


**Figure S10. Changes to zoospore density when vaccination provides both tolerance and resistance.** Generalized Additive Model summary of modeled changes in zoospore density (green-purple color scale) as a function of vaccination-induced increases in (a) anti-infection resistance (decrease in infection establishment), (b) anti-transmission resistance (decrease in pathogen shedding), or (c) anti-growth resistance (increase in pathogen clearance; x-axis) and tolerance (increase in infection induced mortality threshold; y-axis), relative to simulations of an untreated control population. Population coverage was kept at 75% across all simulations. Deeper green shades represent reductions and deeper purples represent increases in zoospore density as compared to unvaccinated populations. Contour lines define increments of 20% change relative to vaccine-free simulations. Zoospore density decreases with increasing levels of resistance, but remains unchanged when resistance is low and tolerance is high.

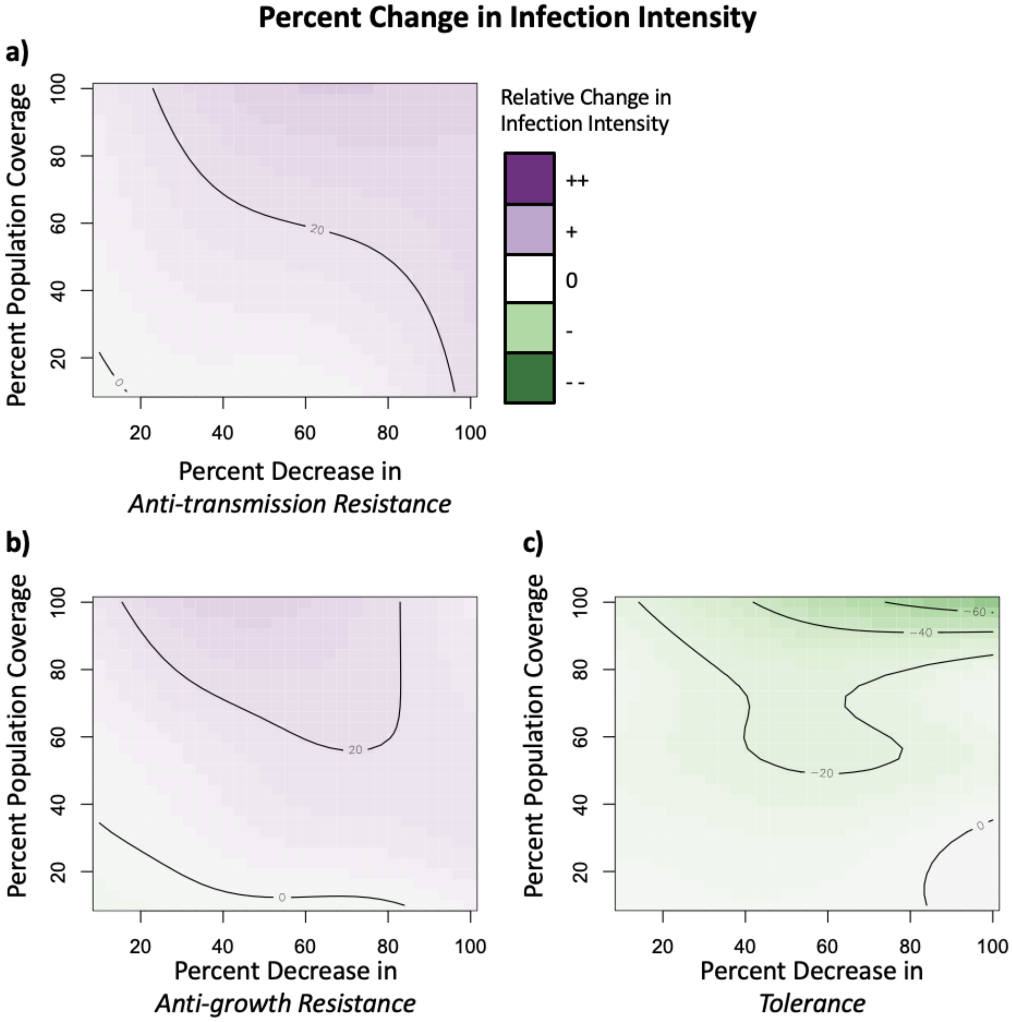


**Figure S11. Changes to infection intensity when vaccination reduces immunity across increasing population coverage.** Generalized Additive Model summary of modeled changes in infection prevalence (green-purple color scale) as a function of vaccination-induced decreases in (a) anti-transmission resistance, (b) anti-growth resistance (decrease in pathogen clearance) or (c) tolerance (decrease in infection induced mortality threshold; x-axis) and increasing population coverage (y-axis), relative to simulations of an untreated control population. Deeper green shades represent reductions and deeper purples represent increases in infection intensity as compared to unvaccinated populations. Contour lines define increments of 20% change relative to vaccine-free simulations. Infection intensities increase with (a, b) high population coverage and decreased levels of resistance, but infection intensities decrease with (c) high population coverage and decreased tolerance.

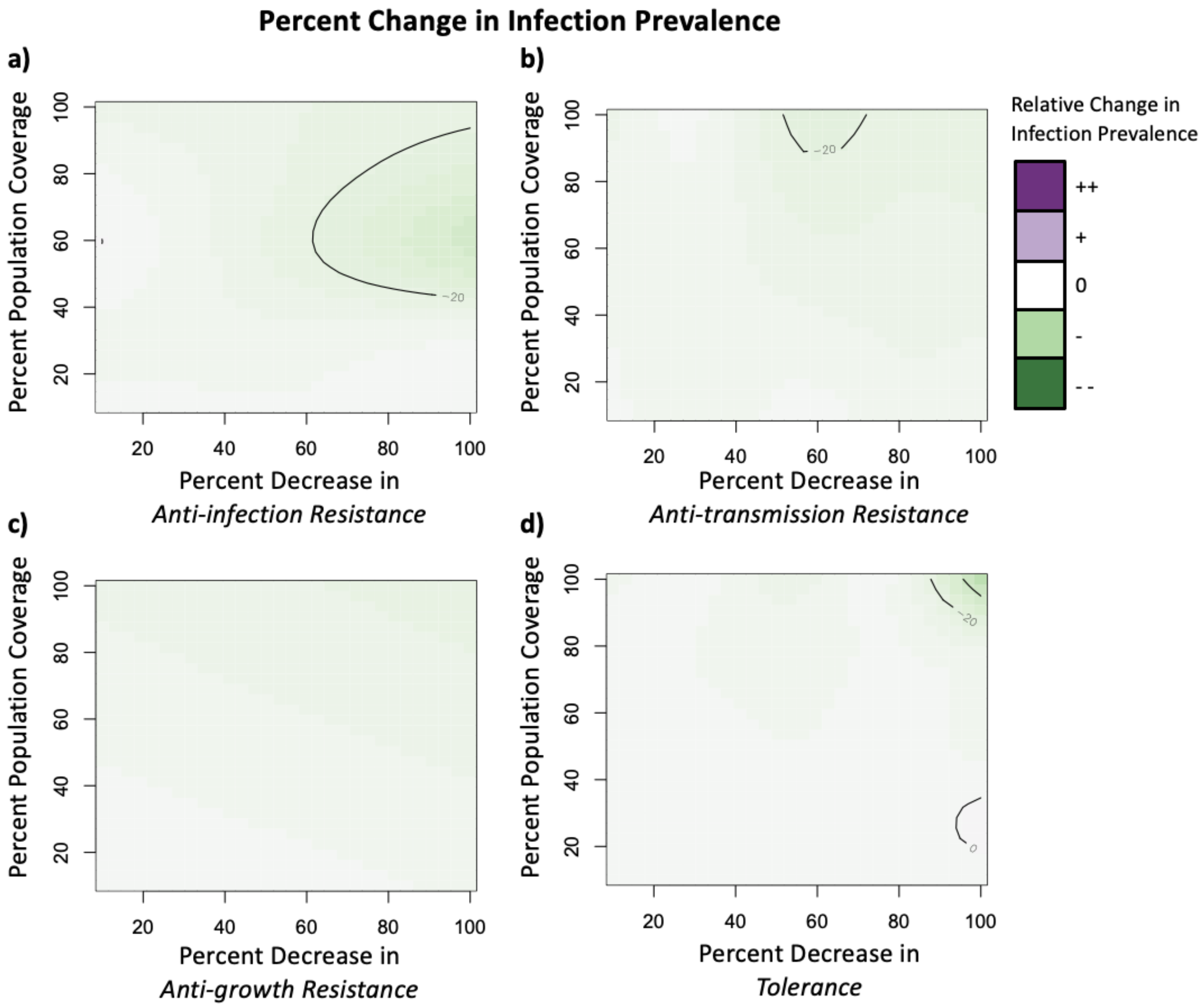


**Figure S12. Changes to infection prevalence when vaccination is harmful, across increasing population coverage.** Generalized Additive Model summary of modeled changes in infection prevalence (green-purple color scale) as a function of vaccination-induced decreases in (a) anti-infection resistance (increase in infection establishment), (b) anti-transmission resistance (increase in pathogen shedding), (c) anti-growth resistance (decrease in pathogen clearance) or (d) tolerance (decrease in infection induced mortality threshold; x-axis) and increasing population coverage (y-axis), relative to simulations of an untreated control population. Deeper green shades represent reductions and deeper purples represent increases in infection prevalence compared to unvaccinated populations. Contour lines define increments of 20% change relative to vaccine-free simulations. Infection prevalence only decreases under certain scenarios of decreasing (a) anti-growth resistance, (b) anti-transmission resistance, and (d) tolerance, but remains unchanged regardless of (c) level of decreased anti-growth resistance.

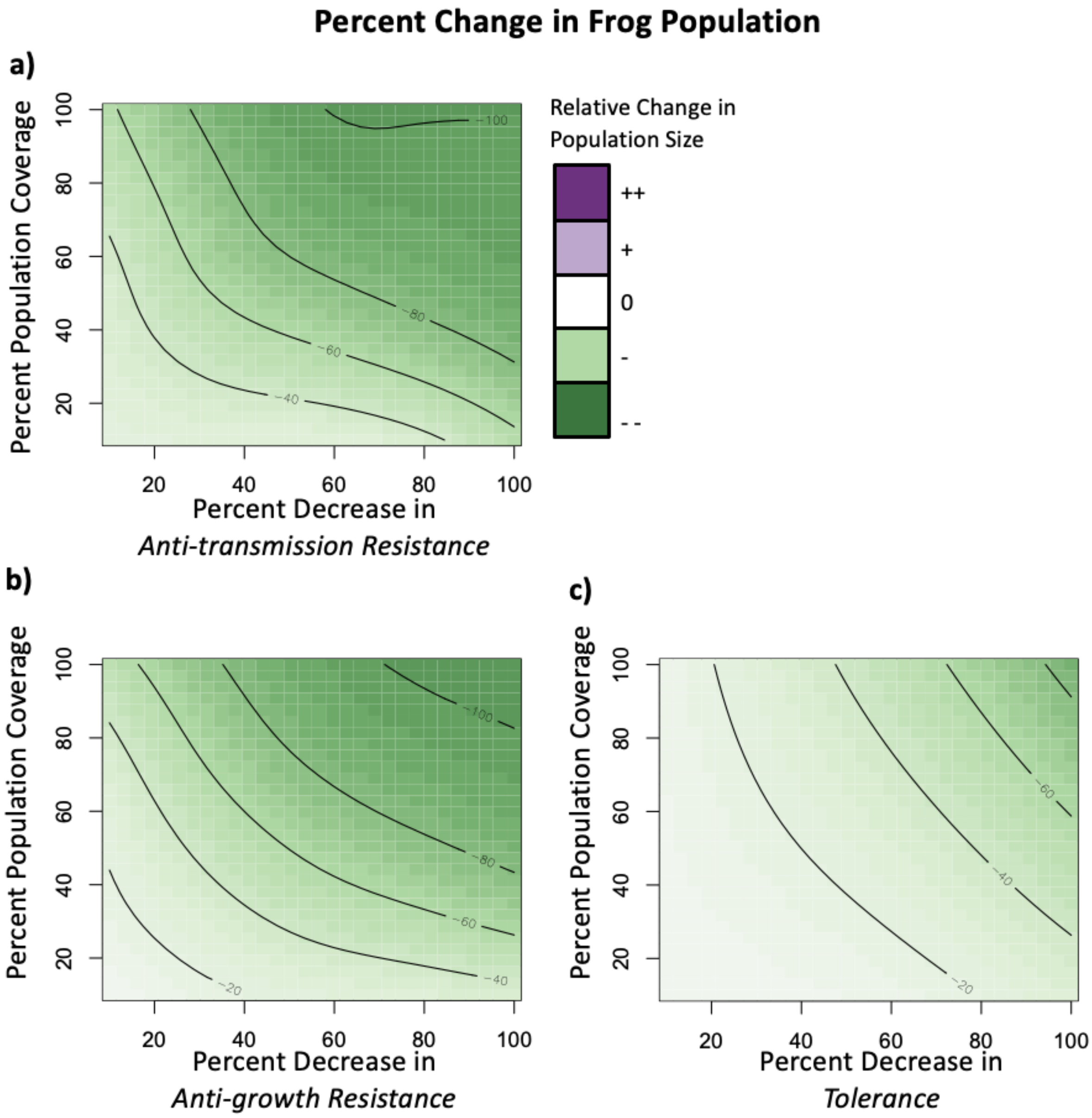


**Figure S13. Changes to frog population size when vaccination is harmful, across increasing population coverage.** Generalized Additive Model summary of modeled changes in infection prevalence (green-purple color scale) as a function of vaccination-induced decreases in (a) anti-transmission resistance, (b) anti-growth resistance (decrease in pathogen clearance) or (c) tolerance (decrease in infection induced mortality threshold; x-axis) and increasing population coverage (y-axis), relative to simulations of an untreated control population. Deeper green shades represent reductions and deeper purples represent increases in population size as compared to unvaccinated populations. Contour lines define increments of 20% change relative to vaccine-free simulations. Frog populations decrease with decreasing levels of resistance or tolerance.

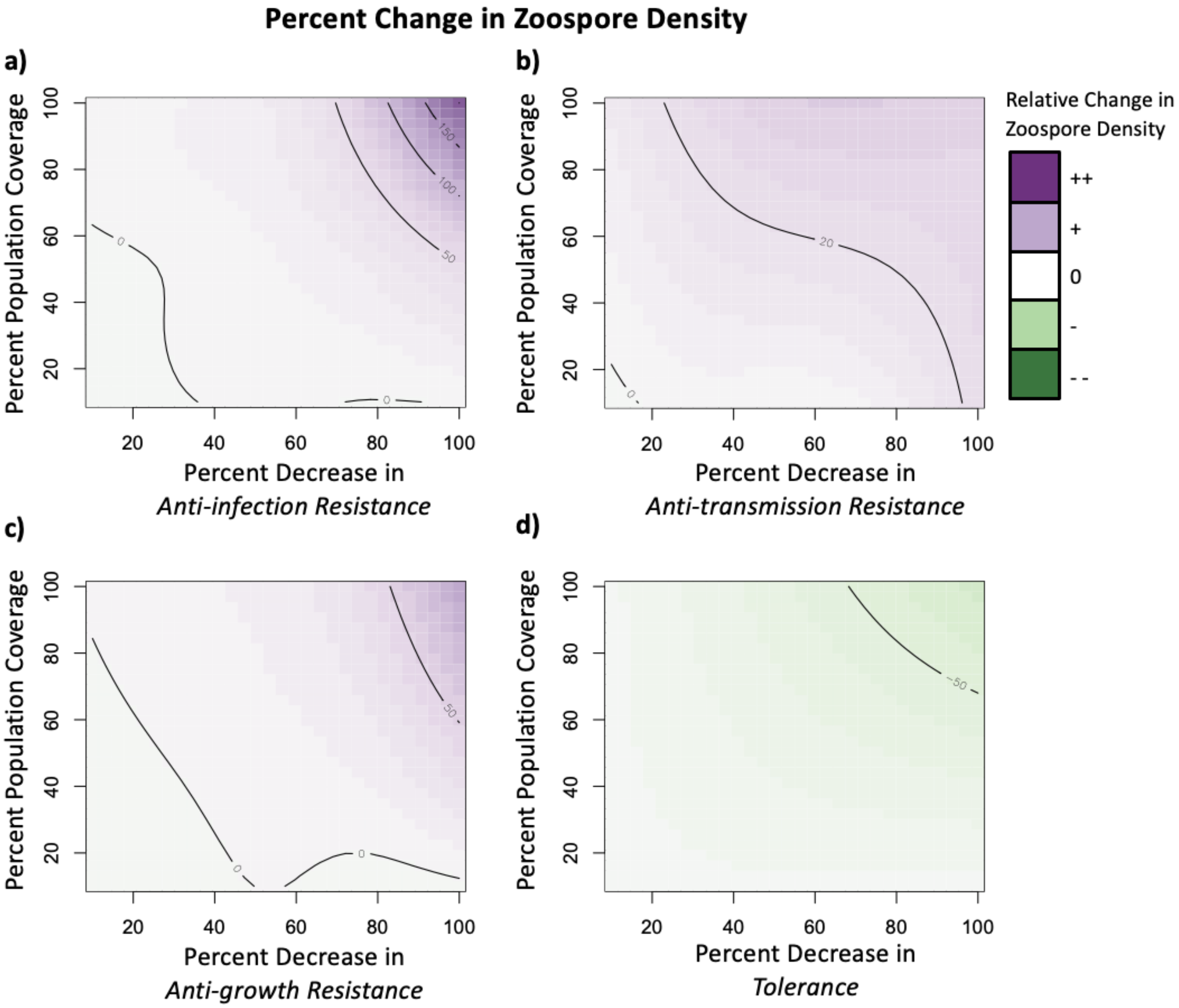


**Figure S14. Changes to zoospore density when vaccination is harmful, across increasing population coverage.** Generalized Additive Model summary of modeled changes in infection prevalence (green-purple color scale) as a function of vaccination-induced decreases in (a) anti-infection resistance (increase in infection establishment), (b) anti-transmission resistance (increase in pathogen shedding), (c) anti-growth resistance (decrease in pathogen clearance) or (d) tolerance (decrease in infection induced mortality threshold; x-axis) and increasing population coverage (y-axis), relative to simulations of an untreated control population. Deeper green shades represent reductions and deeper purples represent increases in zoospore density as compared to unvaccinated populations. Contour lines define increments of 50% change relative to vaccine-free simulations. Zoospore densities increase as resistance decreases (a-c), but zoospore densities decrease with high reductions in tolerance.

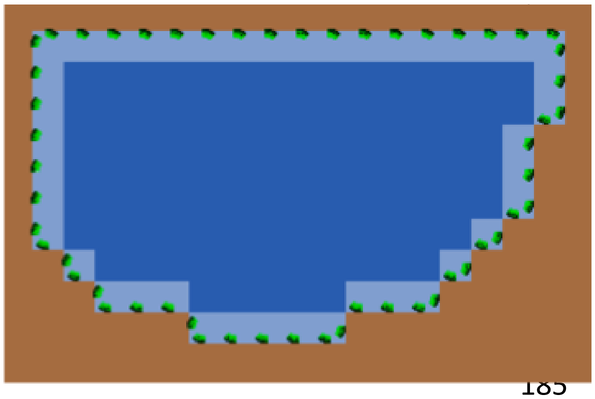

**Figure S15. Netlogo user interface graphical display of the Bd-amphibian-vaccine model's spatial structure.** There are types of environmental patches in this model: 1) perimeter pond patches (light blue), 2) deep pond patches (dark blue), and 3) terrestrial patches (brown). Amphibians are represented as green and black objects.

**Table S1.** Major model processes, defined as the following: amphibian phenology and ecology, implementation of acquired immunity (vaccination), between-host transmission, and within-host infection processes.

| **Major Model Processes** | | | |
|---|---|---|---|
| **Process** | **Baseline parameter** | **Variation in parameter** | **Procedural info** |
| ***Amphibian phenology and ecology*** | | | |
| Within-season dynamics | Last-day = 90 ticks (to match 3 month transmission season from tadpole hatching to metamorphosis). | No | Simulation ends at day 90. |
| Natural tadpole mortality | tad-mort = 0.06 $day^{-1}$ (1) | No | Probability of mortality each time step. |
| Natural metamorph mortality | meta-mort = 0.02 $day^{-1}$<br><br>Determined through pattern-matching (Fig. S16). | No | Probability of mortality each time step. |

| | | | |
|---|---|---|---|
| Tadpole movement | Proportion of tadpole population that moves each time step (‘t-movement’) = 0.25. | No | 25% of tadpoles move to a new perimeter pond patch each time step. |
| Metamorph movement | Proportion of metamorphs on land at each time step (‘m-land’) = 0.1. | No | Metamorphs move to a new patch each time step. 10% of metamorphs are on terrestrial patches and 90% are on perimeter pond patches. |
| Metamorphosis | Tadpoles transition to metamorphs between day 55-74.<br><br>Determined through pattern-matching (Fig. S16). | No | Beginning on day 55, each tadpole has a 11% chance of transitioning to a metamorph. On day 74, all remaining tadpoles become metamorphs. Metamorphs retain infections (2) and all immune traits from tadpole state. |
| ***Acquired immunity*** | | | |
| Vaccination | Host vaccination status is ‘immunized’ = 1 if host is vaccinated or 0 if host is unvaccinated.<br><br>Vaccine constants (c_est, c_clear, c_shedding, and c_smax) given vaccination status are specified per scenario.<br><br>v-efficacy = 1 relative_variation = 10%. | To allow for a range of individual variation in response vaccination, we allow for +/- 10% differences in response to the baseline vaccine efficacies specified in each scenario. | We use constants to modulate baseline parameters regarding infection establishment (‘c_est’), infection clearance (‘c_clear’), infection shedding (‘c_shedding’), and infection-induced mortality (‘c_smax’).<br><br>Vaccine parameter c_est, c_clear, c_shedding, and c_smax serve as exponential modulating factors (exponential to prevent values from becoming negative) that adjust the baseline values for probability of successful infection establishment given zoospore exposure, probability of zoosporangia clearance, zoospores shed per |

| | | | |
|---|---|---|---|
| | | | zoosporangia, or zoosporangia threshold above which mortality occurs, respectively, given a host's vaccination status and 'imm' parameter. Positive values of constants increase baseline parameters, while negative values decrease baseline parameters, and constants = 0 when hosts are unvaccinated or vaccination does not impact a given process. |
| Vaccination coverage | Specified per scenario. | No | On day of vaccination (default = day 0), the specified proportion (70% coverage = 0.7) of the tadpole population is randomly selected. Immune parameters of selected uninfected tadpoles will be adjusted according to the scenario's vaccine efficacy (see "vaccination" process). Immune parameters of selected but infected tadpoles will remain unchanged given that Bd-metabolites have been found to be ineffective at inducing acquired resistance in frogs previously challenged with Bd (3). |
| ***Between-host infection processes*** | | | |
| Host exposure to zoospores | amount of environmental units each host is exposed to per day = 0.25 (unitless).<br><br>Determined through pattern-matching (Fig. S17). | No | Transmission is determined by draws from a multinomial probability distribution where each host's likelihood of infection with a zoospore is determined by its establishment and exposure parameter. (4) |
| Infection Establishment | probability of successful infection establishment upon exposure ('est') = 25%. | Varies depending on host vaccination status and vaccine efficacy scenario. | |

| | | | |
|---|---|---|---|
| Zoospore removal from pool | zoospore mortality factor (‘z-mort’) = 2 $day^{-1}$ (5). | No | Zoospores are removed from patches as a function of exposure to hosts and exponential decay determined by the mortality factor. |
| ***Within-host infection*** | | | |
| Zoosporangium maturation | NA | No | Upon a zoospore successfully establishing an infection, it moves between one of 4 prezoosporangia stages (pz0-pz4) per day to approximate the 4 days it takes for a zoosporangium to mature before shedding (6). |
| Infection clearance | baseline_spn_clearance = 0.20 $day^{-1}$ (7). | Varies depending on host vaccination status and vaccine efficacy scenario. | Binomial draw from probability of zoosporangia clearance to determine how many sporangia survive on each host. |
| Zoosporangia shedding | 17.8 zoospores produced per zoosporangium per day (7). | Varies depending on host vaccination status, ‘imm’ value, and c_shedding. | Zoosporangia shed 40% of zoospores to the patch the host is currently on, 50% of zoospores to a neighboring perimeter or inner patch, and 10% of zoospores directly re-expose the host that produced them. |
| Self-reinfection | Proportion of zoospores that a host releases and is re-exposed to = 0.1 (7). | Varies depending on host vaccination status, ‘imm’ value, and c_est. | Zoospores that reinfect host is calculated by multiplying the number of zoospores that host releases with the probability of infection given establishment (‘est’) |
| Infection-induced mortality | Threshold of sporangia above which mortality occurs (‘smax’) = 562.<br><br>Derived from (7) which cites maximum 10,000 zoospores released per day. We divided 10,000 zoospores by 17.8 zoospores shed per | Varies depending on host vaccination status, ‘imm’ value, and c_smax. | Metamorph dies if the zoosporangia load reaches threshold value (‘smax’). |

| | | | |
|---|---|---|---|
| | zoospore per day to approximate 562 as the maximum number of zoosporangia per host. | | |
| Tadpole sporangia carrying capacity | Maximum zoosporangia burden for a tadpole ('s_k') = 10,000.<br><br>Determined through pattern-matching (Fig. S17). | No | If a tadpole has 10,000 zoosporangia, it cannot be further infected. However, it can continue to be exposed to zoospores. |

***<u>SI References</u>***
